# Benchmarking algorithms for RNA velocity inference

**DOI:** 10.64898/2026.01.03.697314

**Authors:** Kexin Huang, Yu Zhou, Tiangang Wang, Xiao Li, Xinlong Zhao, Xi Liu, Liyu Huang, Xiaobo Zhou, Jiajia Liu

**Author notes:** These authors contribute equally to this work (Kexin Huang, Yu Zhou, Tiangang Wang). Address correspondence to: Jiajia Liu, Ph.D. McWilliams School of Biomedical Informatics The University of Texas Health Science Center at Houston 7000 Fannin St., Houston, TX 77030 Xiaobo Zhou, Ph.D. McWilliams School of Biomedical Informatics The University of Texas Health Science Center at Houston 7000 Fannin St., Houston, TX 77030.

## Abstract

RNA velocity is a computational framework for single-cell RNA sequencing (scRNA-seq) that estimates the future transcriptional state of individual cells, thereby capturing the direction and rate of cell state transitions rather than providing a purely static snapshot. Since its introduction in 2018, multiple RNA velocity methods have been developed, differing in their modeling assumptions, required inputs, computational complexity, and robustness. However, there remains limited consensus on how best to evaluate these methods or on which tools are most reliable under specific biological and technical settings. Here, we perform a systematic comparison of 29 velocity inference algorithms across 114 simulated datasets with known ground-truth cell dynamics and 62 real scRNA-seq datasets, and we extend the evaluation to spatial and multi-omics levels where velocity is increasingly applied. We benchmark RNA velocity methods using a unified framework that decomposes performance into four practical dimensions: accuracy, scalability, stability, and usability. Our results show that performance rankings vary substantially across metrics and datasets, indicating no single method is uniformly optimal and that practical deployment is often constrained by feasibility and robustness as much as by accuracy. Based on these results, we provide actionable guidance for selecting RNA velocity tools according to data modality, available priors, and computational constraints. Finally, we identify key bottlenecks that currently limit RNA velocity development and deployment, including scalability to large size of datasets, sensitivity to gene selection, and the lack of genuinely multimodal and spatially explicit velocity models for spot-based technologies.

## Introduction

The rapid advance of high-throughput sequencing, especially single-cell RNA-seq and spatial omics, has opened new avenues for studying cellular dynamics such as differentiation, heterogeneity, and the cell cycle ^1,2^. Traditional trajectory inference approaches reconstruct these dynamics by imposing predefined structural assumptions, such as linear, tree structure, or disconnected graph-based topologies ^3^. These trajectory inference methods are onto static snapshots of gene expression, often requiring prior biological knowledge to specify developmental origins and endpoints ^4^. RNA velocity resolves this limitation by inferring the direction and rate of transcriptional change directly from scRNA-seq, modeling the balance between unspliced precursor transcripts and their spliced mature counterparts to predict each cell’s future state ^5^. Because velocity is estimated locally and independently of any global trajectory model, it does not rely on a predefined lineage structure and remains agnostic to the underlying topology of cell state space ^4^. As a result, RNA velocity accommodates diverse trajectory topologies including linear paths, bifurcations, cycles, and more complex graphs. Recent work extends the concept beyond transcript abundance through multimodal integration with chromatin accessibility, joint RNA-protein measurements, and metabolic labeling that provides temporal ground truth ^6,7^. In spatial transcriptomics, velocity vectors can be mapped onto tissue coordinates to reveal pattern formation, morphogenetic flows, neuronal migration, and tumor-stroma interactions in situ ^8,9^. As a result, RNA velocity is used not only to chart development and lineage commitment but also to interrogate disease progression, immune activation and exhaustion, cell-cycle dynamics, tissue regeneration, and treatment response, adding a time-aware layer to single-cell state maps ^10,11^.

Building on this foundation, a wide array of velocity inference methods has appeared across modalities and is now woven into end-to-end single-cell analysis. In several published pipelines, including bollito, CellexalVR, and “awesome-single-cell” collections (https://github.com/seandavi/awesome-single-cell), RNA velocity is treated as one of core analysis layers ^12,13^. While RNA velocity methods vary in algorithms, priors, and outputs, two features most clearly distinguish them: the kinetic paradigm they adopt or learn, ranging from steady-state assumptions to full dynamical inference, and whether they merely project local directions on a fixed embedding or reconstruct a global vector field that captures branching and other complex trajectory topologies ^5^. Early velocity tools relied on a steady-state assumption, estimating direction from spliced-unspliced balances, they were efficient but sensitive to normalization and noise, and often faltered with transient kinetics or complex branching ^14^. Newer methods relax the steady-state constraint with dynamical models, reconstruct smoother vector fields and fate probabilities, and integrate multimodal or spatial data, providing more robust directionality and richer downstream inference ^15^.

Given the diversity in velocity inference methods, it is essential to quantitatively assess their accuracy, scalability, robustness and usability. While there are limited attempts at only a small number of benchmarking studies have tacking this issue and they lack of a comprehensive comparison of velocity inference methods based on a large number of datasets with different biological contexts, trajectory topologies and sample sizes. This gap poses a significant challenge for new users, who are confronted with an overwhelming choice of velocity inference methods, without a clear guidance on which approach is best suited to their specific analytical needs. Moreover, a systematic assessment of the strengths and limitations of existing methods is necessary to guide future methodological development and to identify key areas for improving the current state-of-the-art.

Thus, in this study, we benchmarked 29 velocity inference methods, including 20 RNA velocity inference methods ^9,14–32^, 7 multi-omics velocity inference methods^6,7,9,31–34^, and 2 velocity-based cell cycle inference methods ^35,36^ (**Supplementary Table S1**) across 176 single-cell datasets (**Supplementary Tables S3, S4**). The performance was evaluated using 17 metrics across four complementary dimensions: accuracy, stability, scalability, and usability (**Fig. 1**). We observed substantial complementarity among existing velocity methods, with different approaches performing optimally depending on the characteristics of the data. To facilitate practical method selection, we developed a public knowledgebase of our benchmarking results and guidelines (https://relab.xidian.edu.cn/RNAVelocity/), providing detailed performance of each method and their comparisons. Finally, our results highlighted some challenges for current methods, and our evaluation strategy can be useful to spearhead the development of new tools that accurately infer velocity on ever more complex use cases.

**Fig 1.**
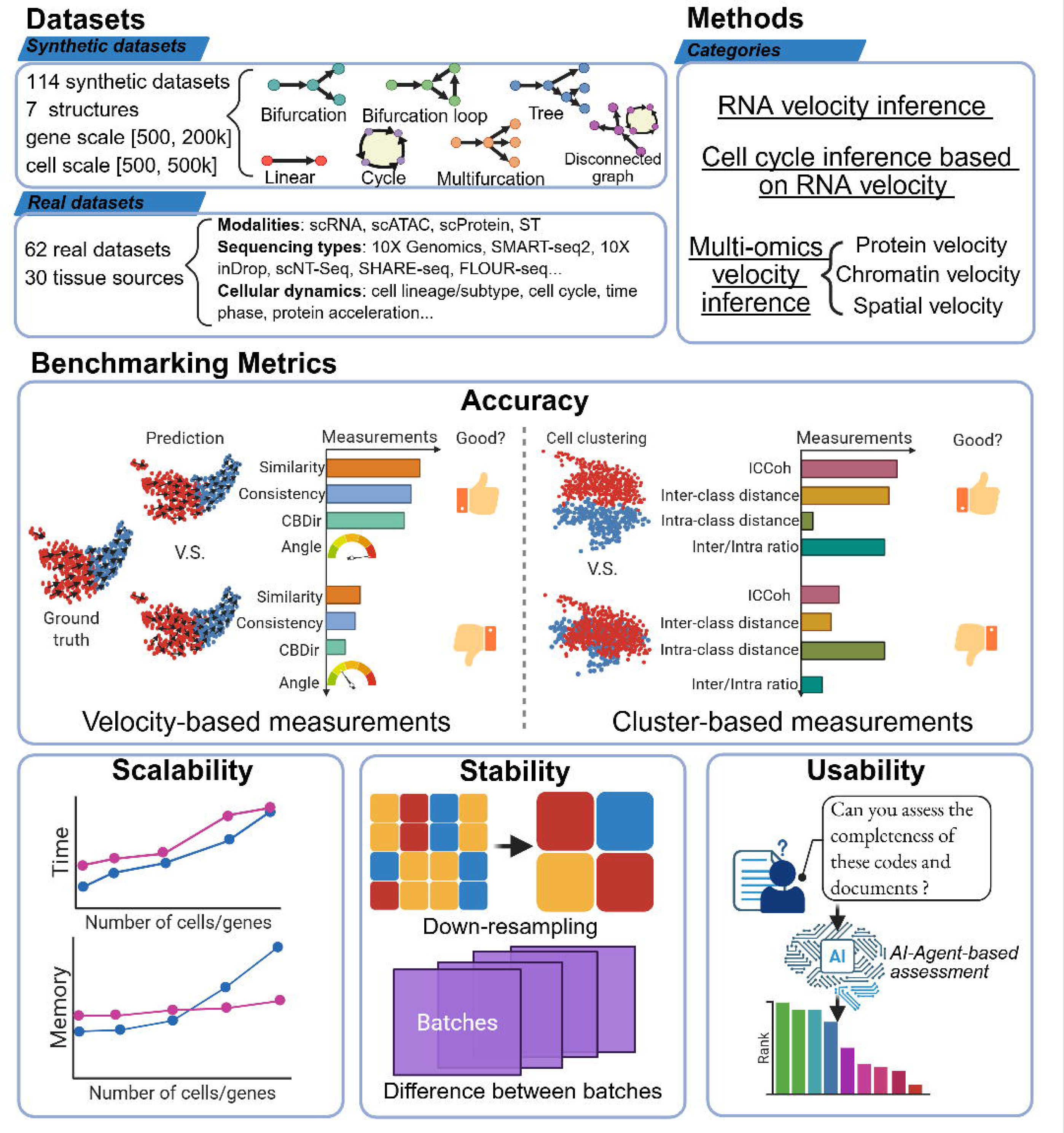
Overview of datasets, method categories, and benchmarking metrics for velocity inference. Top left, dataset compendium used in this study. Top right, benchmarked method categories, including RNA velocity inference, cell-cycle inference based on RNA velocity, and multi-omics velocity inference (protein velocity, chromatin velocity, and spatial velocity). Middle, accuracy metrics grouped into velocity-based measurements and cluster-based measurements. Bottom, additional evaluation dimensions.

## Results

### Velocity inference methods and evaluation framework

We benchmarked a broad panel of velocity inference methods across diverse biological and technical settings to provide a decision-oriented assessment of their reliability and practical usability. The methods vary in requirements, including implementation platform, input format, and any additional inputs except standard spliced and unspliced counts (**Fig. 2**). They also span a wide range of dynamical assumptions, from near–steady-state or locally linear models to explicit kinetic, probabilistic, and deep generative approaches, and differ in output targets, including cell-wise velocity vectors, cell-to-cell transition graphs or probabilities, and latent-time fields.

**Fig 2.**
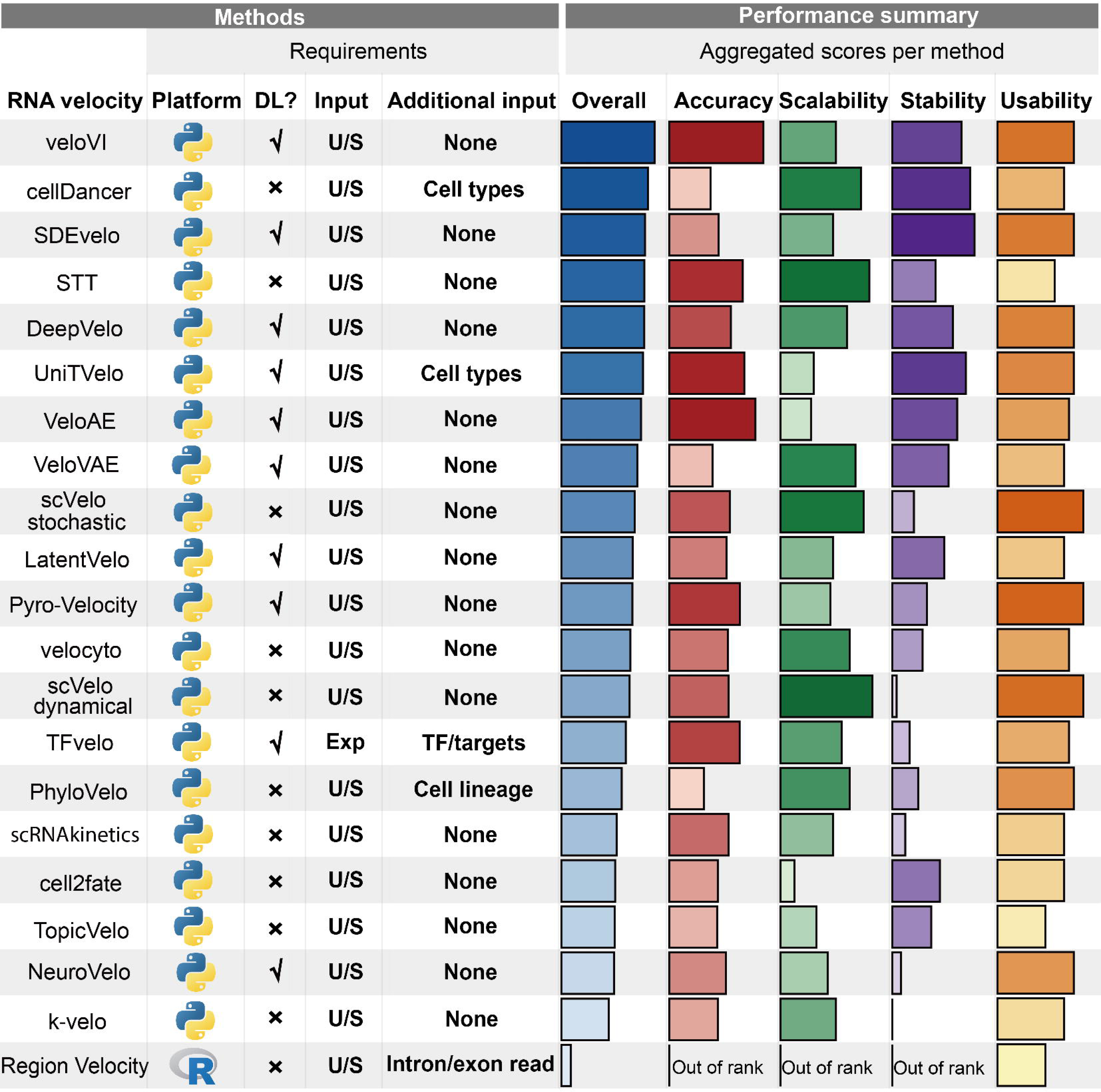
Summary of method requirements and aggregated benchmarking performance for RNA velocity inference. Left, practical requirements for each evaluated RNA velocity method. Right, aggregated performance summaries across the full benchmark, reported as an overall score and four evaluation dimensions. The overall score is the mean of metric-specific ranks pooled across both simulated and real datasets, with higher ranks indicating better overall performance. Methods that could not be meaningfully ranked for a dimension are marked as out of rank.

These differences affect robustness under distinct dynamical scenarios. To account for this, we first benchmarked methods across a set of controlled simulation settings with different dynamical structures and scale regimes. The dynamic structures cover linear, bifurcating, cycle, trifurcating, bifurcating loop, consecutive bifurcating and disconnected structures. This setup mirrors the rationale used in trajectory inference benchmarking, where topology is the key discriminant regard of cell dynamics. We then evaluated performance on 62 real single-cell datasets, encompassing processes such as differentiation, cell-cycle dynamics, disease progression, and immune activation, to assess practical utility in realistic scenarios.

Methods were evaluated along four complementary dimensions: (1) Accuracy, quantified using metrics for vector-level agreement, cluster coherence, and downstream biological consistency, with additional assessment of spatial coherence for spatial velocity outputs (see Methods). (2) Scalability, measured via Docker-recorded runtime and memory usage, complemented by peak CPU and GPU usage to capture transient resource bursts. (3) Stability, assessed via robustness to input perturbations through systematic down-sampling and multi-run reproducibility, particularly relevant for deep learning-based methods. (4) Usability, summarized by an aggregated score reflecting installation friction, documentation quality, maintainability, and user-facing diagnostics. An overall performance score was defined as the mean of metric-specific ranks aggregated across both simulated and real datasets, with higher ranks indicating better performance (**Fig. 2**).

Across this aggregated summary, veloVI ^17^ ranked first overall, followed by cellDancer ^18^, SDEvelo ^31^, STT ^32^, and DeepVelo ^22^, with UniTVelo ^19^ and VeloAE ^27^ also placing in the upper tier. Importantly, the top-ranked methods were not those that dominated a single metric, but those that avoided severe weaknesses across accuracy, scalability, stability, and usability, yielding comparatively balanced profiles. We will discuss each evaluation criterion in detail (**Fig. 3A**), after which we conclude with guidelines for method users and future perspectives for method developers.

**Fig 3.**
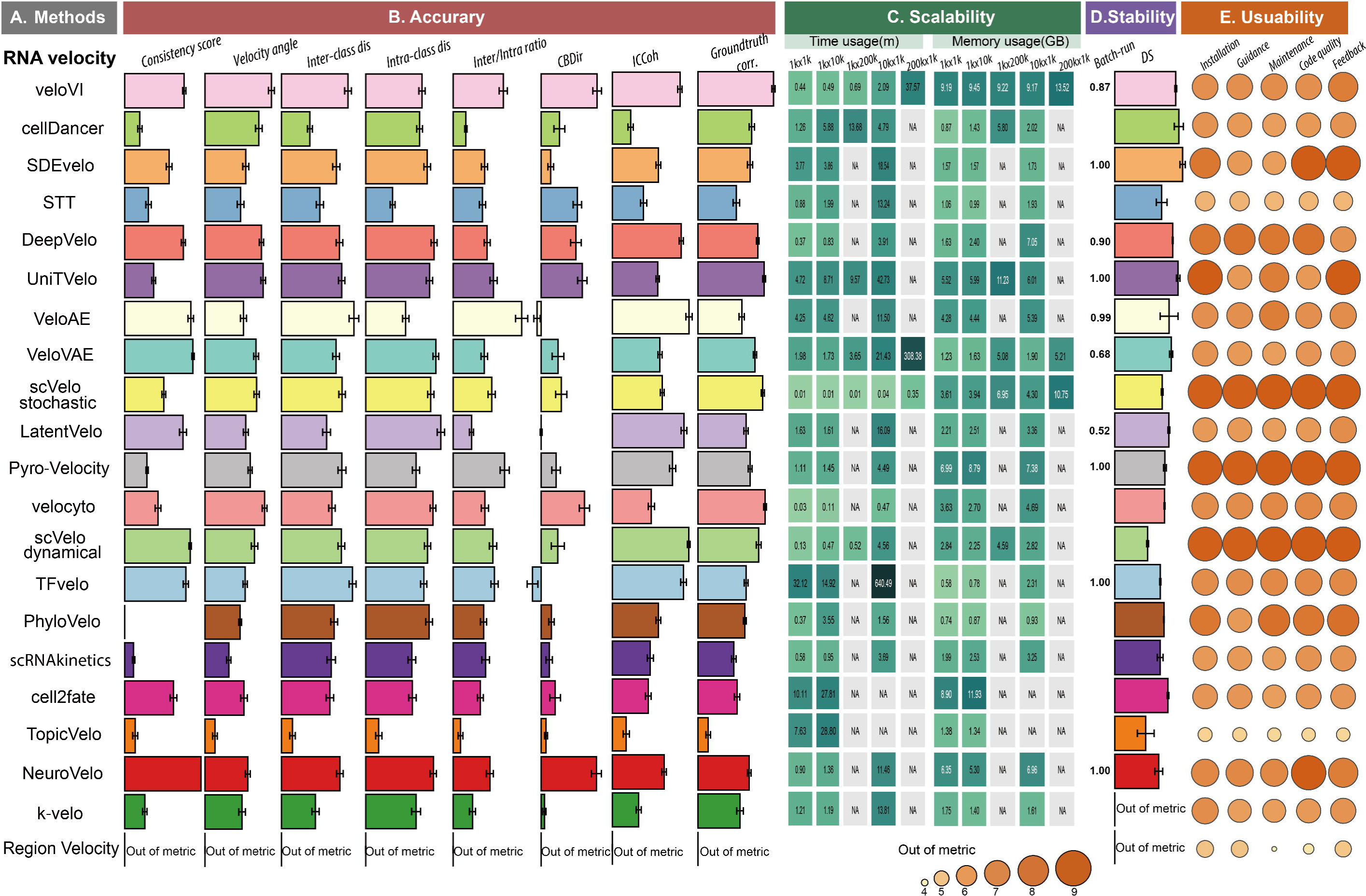
Metric level benchmarking results across accuracy, scalability, stability, and usability based on simulated datsets. **A**, Methods. Evaluated RNA velocity inference tools, ordered consistently across panels. **B,** Accuracy. Method performance on eight complementary accuracy metrics used for simulated datasets. Longer bars indicate better performance for that metric after orienting scores so that higher is better. **C,** Scalability. Runtime and memory usage measured across increasing dataset sizes for each method. Cells show the observed time and memory values for each size setting, with NA indicating that the run did not complete or was not available under the standardized pipeline. **D,** Stability. Reproducibility and robustness summarized by two tests, multi run agreement across repeated runs and down sampling sensitivity under controlled removal of cells or genes. Higher values indicate more stable results. **E,** Usability. Practical usability scores assessed across installation, guidance quality, maintenance, code quality, and user facing feedback, where larger circles indicate higher scores. Methods marked as out of metric were not evaluated for that panel under the unified framework.

### Accuracy

Accuracy was evaluated using eight complementary metrics that can be broadly categorized into velocity-based measurements and clustering-based measurements, capturing both local dynamical correctness and global structural preservation (**Fig. 3B**, **see Methods**). Velocity-based measurements assess the correctness and coherence of inferred velocity vectors, including ground-truth correlation, velocity angle consistency, cross-boundary direction correctness (CBDir) ^27^ and in-cluster coherence (ICCoh) ^27^ and consistency score ^16^. Ground-truth correlation, applied only to simulated datasets, directly quantifies alignment between inferred and true velocity vectors. Velocity angle consistency evaluates whether inferred directions follow the expected lineage orientation in the embedding. CBDir assesses cross-boundary directionality by testing whether velocities at cluster interfaces point from putative source states toward target states ^27^. ICCoh measures within-cluster directional coherence, while the consistency score and its peak statistic quantify local neighborhood smoothness and whether high coherence is broadly supported across cells ^27^. Clustering-based measurements were used to evaluate whether velocity inference preserves the global organization of cell states. Inter-class distance, intra-class distance, and their inter-intra ratio summarize state separation and compactness, capturing how well velocity-associated structure maintains clustering geometry and global topology in the cell-state space ^20^.

Across metrics and datasets, method rankings differed markedly, indicating that no single approach is uniformly optimal across all accuracy criteria. This variability is expected because velocity methods differ in their core dynamical assumptions and inferential targets, for example, reliance on steady-state approximations versus explicit dynamical modeling, treatment of kinetic parameters as cell-agnostic versus state dependent, and whether outputs take the form of denoised vector fields, probabilistic latent-time models, or transition operators with uncertainty quantification. At the aggregate accuracy level summarized in **Fig. 2**, the highest accuracy ranks were achieved by methods that explicitly couple kinetic modeling with representation learning or probabilistic inference, including veloVI, VeloAE, UniTVelo, STT, Pyro-Velocity, TFvelo, and DeepVelo. Despite substantial differences in implementation, these methods share a common theme that they reduce sensitivity to sparsity and model misspecification by either learning structured latent representations of dynamics or by introducing probabilistic structure that shares information across cells and genes and exposes uncertainty, rather than relying solely on per-gene steady-state deviations.

Different accuracy metrics consistently highlighted different subsets of velocity inference methods, reflecting how modeling choices translate into distinct strengths under particular evaluation criteria. For ground-truth correlation, methods that explicitly learn flexible continuous-time dynamics, such as NeuroVelo, ranked highest, consistent with their ability to recover non-linear velocities without relying on steady-state assumptions. Velocity angle consistency and CBDir emphasize correct global orientation and cross-boundary directionality. Unified-time and deep dynamical models, including UniTVelo, probabilistic latent-time frameworks such as veloVI, and continuous-time learners such as DeepVelo, achieve high scores on these metrics because they jointly infer latent time and kinetics, thereby stabilizing directionality and reducing spurious reversals across state boundaries. For local coherence-driven metrics (ICCoh and consistency score), representation-learning and probabilistic frameworks, including VeloAE, veloVI, and Pyro-Velocity, consistently performed well, reflecting their denoised and manifold-consistent velocity fields enabled by information sharing across genes and cells. Finally, three clustering-based metrics assessing global structure preservation highlighted veloVI, STT, and UniTVelo, which combine coherent within-state dynamics with well-resolved transitions between states.

In summary, the metric-by-metric dispersion indicates that accuracy in RNA velocity is not a single axis but a bundle of partially competing desiderata. In practice, method choice should therefore be tied to the intended downstream use. Workflows that prioritize reliable lineage orientation and boundary transitions should weight CBDir and velocity-angle style criteria more heavily, whereas analyses centered on vector-field visualization and local dynamical interpretation should emphasize ICCoh and consistency-style coherence metrics. The aggregate accuracy rankings in **Fig. 2**. provide a useful navigation index, but the per-metric breakdown in **Fig. 3B** and **Supplementary Fig. 1** is necessary to understand which property each top-ranked method is actually optimizing and which failure modes remain plausible for a given biological question.

### Scalability

We assessed computational scalability using Docker-recorded runtime and memory usage for 21 RNA velocity methods across synthetic datasets with different sizes of genes and cells, complemented by peak resource occupancy during execution (**Fig. 3C**). All Docker environments were deployed on two high-performance servers, including a GPU-enabled system for deep learning–based methods and a CPU-focused system for conventional pipelines (**see Methods**). Overall, scalability constraints manifested first as failures to complete on large datasets: across the 100,000 cells ×1,000 genes settings under multiple topologies, only 9-12 out of 21 RNA-velocity methods returned numeric runtimes, and this fraction dropped further for extra-large datasets, where only 3/21 methods completed at 200,000 cells ×1,000 genes and only 2/21 at 500,000 cells ×1,000 genes. Even among methods that did complete, runtime varied dramatically, ranging from less than a second (0.68s for scVelo stochastic) to over an hour (3821.42s for TopicVelo), indicating substantial practical differences in throughput even before accounting for failures. Notably, only a small subset of methods, including scVelo stochastic, veloVI, and VeloVAE, remained usable at 200,000 cells ×1,000 genes, and only veloVI and VeloVAE completed at 500,000 cells ×1,000 genes (**Supplementary Table S5**).

For most methods, runtime was driven mainly by the number of cells rather than the number of genes, with computation time increasing rapidly as cell counts grew. For example, in the bifurcating structure with 1,000 genes, scaling from 1,000 to 100,000 cells increased runtime by approximately 188× for scVelo stochastic (0.205s to 38.548s) and 153× for DeepVelo (16.24s to 2488.63s), consistent with near-linear to mildly superlinear growth over this range. In contrast, for some implementations, gene dimensionality emerged as the dominant bottleneck, leading to sharp slowdowns even at moderate cell numbers. When holding 1,000 cells fixed and increasing genes from 1,000 to 100,000, runtime inflation was highly method-dependent: velocyto increased by ∼512× (1.436s to 736.084s) and PhyloVelo by ∼187× (20.278s to 3786.160s), whereas veloVI remained nearly unchanged (∼1.05×, 26.480s to 27.872s), highlighting distinct algorithmic bottlenecks (**Supplementary Table S5**).

Memory usage was generally moderate among completed runs but highly heterogeneous across methods (**Fig. 3C**). Median Docker-reported memory ranged from 0.70 to 12.02 GB, although this summary is affected by survivorship bias because many method–dataset combinations failed to complete, often due to memory exhaustion. Methods with high baseline memory demand, most notably cell2fate, had the highest median Docker memory in our summary at 12.02 GB and showed frequent non-completion already at moderate scales (≥10,000 cells), indicating limited scalability under typical resource constraints. Several deep-learning pipelines also exhibited sensitivity to memory limits; for example, UniTVelo failed due to explicit memory allocation errors at extra-large dataset sizes. Considering completion at extreme cell counts together with runtime and resource usage, veloVI was as the most computationally robust method for large datasets; VeloVAE can complete at the largest scales but at substantially higher runtime cost.

For the two cell-cycle velocity methods, VeloCycle and DeepCycle, we evaluated scalability using the same Docker environmenrs. VeloCycle was consistently more efficient than DeepCycle in both runtime and memory. Specifically, VeloCycle achieved a lower median runtime of 31.20s compared with 74.66s for DeepCycle, and required substantially less memory, with a median memory of 0.23GB versus 9.00 GB for DeepCycle. These differences were also reflected in peak resource occupancy: VeloCycle exhibited a median CPU peak of 2.04 GB, whereas DeepCycle reached 56.70 GB (**Supplementary Table S6**).

### Stability

Beyond accuracy and scalability, we evaluated stability by assessing robustness to data down-sampling and reproducibility across repeated runs (**Fig. 3D**). To quantify robustness to input perturbations, we down-sampled both cells and genes from a bifurcating simulated dataset containing 1,000 cells and 10,000 genes to 60%, 70%, 80%, 90%, and 95% of the original data, and measured agreement between inferred and ground-truth velocities. Based on the aggregated stability score, the top performers were SDEvelo (0.64), cellDancer (0.60), UniTVelo (0.59), and veloVI and VeloAE (both 0.57), followed by DeepVelo (0.54). Importantly, robustness was not only reflected in the average performance but also in sensitivity to perturbation severity. DeepVelo showed minimal variation across both cell and gene down-sampling, with cell-subsampling scores ranging from 0.53 to 0.54 and gene-subsampling scores ranging from 0.54 to 0.56. UniTVelo and veloVI also exhibited relatively constrained variation under cell down-sampling, with ranges of 0.62 to 0.66 for UniTVelo and 0.55 to 0.59 for veloVI. In contrast, several methods were sensitive to specific perturbations. cellDancer cellDancer showed pronounced variability under cell down-sampling, with scores spanning 0.42 to 0.79. While SDEvelo was more affected by gene down-sampling, with scores ranging from 0.44 to 0.71 (**Fig. 3D**, right column).

For deep learning-based RNA velocity methods that involve stochastic training or optimization, we further assessed multi-run reproducibility to evaluate the impact of random initialization and training stochasticity. Each deep learning-based method was run five times on one real dataset and one simulated dataset, and reproducibility was quantified using the mean pairwise correlation across runs. Among the ten methods with evaluable repeated-run outputs, NeuroVelo, Pyro-Velocity, SDEvelo, TFvelo, and UniTVelo showed near-perfect reproducibility, each achieving an average reproducibility score of 1.00, with VeloAE also exhibiting high reproducibility (0.993). In contrast, DeepVelo and veloVI were reproducible but clearly less stable across repeats, with average scores of 0.903 and 0.871, respectively, while VeloVAE and LatentVelo showed substantially lower repeatability at 0.678 and 0.521 (**Fig. 3D**, left column). Taken together, these results indicate that down-sampling robustness and run-to-run reproducibility vary independently across tools, and that for some stochastic methods, reliable use may require either fixing random seeds or aggregating results across repeated runs.

### Usability

To quantify practical usability, we summarized each tool with an overall usability index (range from 0 to10, higher is better) that integrates installation friction, usage guidance, maintenance signals, code quality, and the clarity of user feedback during execution from tool websites and their open-source repositories (**Supplementary Table S7**). Among the 20 evaluated RNA-velocity tools, usability scores ranged from 5.0 to 9.0 (median 7.5; mean 7.33), indicating that most methods are readily usable but differ substantially in day-to-day overhead (**Fig. 3E**). Mature and widely adopted toolkits such as scVelo and Pyro-Velocity achieved the highest usability (both 9.0/10), reflecting streamlined installation, comprehensive documentation and user-friendly examples, and active community with issue tracking. A second tier of tools, including PhyloVelo, UniTVelo, SDEvelo, veloVI, NeuroVelo, DeepVelo, scored 8.0/10, offering runnable notebooks and reasonable guidance, though requiring heavier modeling stacks or more careful configuration. We occasionally encountered minor errors during execution these six tools, but the issues were generally straightforward to diagnose and resolve. In contrast, more niche or less user-facing implementations scored lower (e.g., TopicVelo and Region Velocity, both 5.0/10) but for distinct reasons: Region Velocity was mainly constrained by incomplete or insufficient preprocessing guidance, whereas TopicVelo frequently triggered non-trivial runtime errors that were difficult to diagnose and resolve in our benchmarking environment. STT showed moderate usability (6.0/10), primarily due to installation difficulties driven by environment and dependency mismatches. Notably, all surveyed tools are open source and provide sample code, but differences in packaging, dependency management, versioning, and runtime diagnostics translate into non-trivial reproducibility and adoption costs when benchmarking across diverse datasets.

### Performance of cell-cycle velocity inference tools

DeepCycle and VeloCycle are specifically designed for RNA velocity inference along the cell cycle. We compared their performance using both synthetic cyclic data and real cell cycle data, evaluating predictive accuracy, phase composition, gene expression patterns along inferred pseudotime, and computational scalability. Overall, both methods successfully capture cell cycle dynamics, but DeepCycle provides more accurate classification of cell cycle stages, whereas VeloCycle is more scalable and stable on larger datasets.

Across seven clustering metrics, Adjusted Rand Index (ARI), Normalized Mutual Information (NMI), Rand Index (RI), Accuracy, Precision, Recall, and F1 score, DeepCycle consistently outperforms VeloCycle (**Fig. 4A**), indicating better concordance with the reference phase annotations across evaluations. VeloCycle performs comparably but with slightly lower medians across most metrics, suggesting that it prioritizes a lighter-weight inference that preserves overall phase structure without maximizing discrete label assignment.

**Fig 4.**
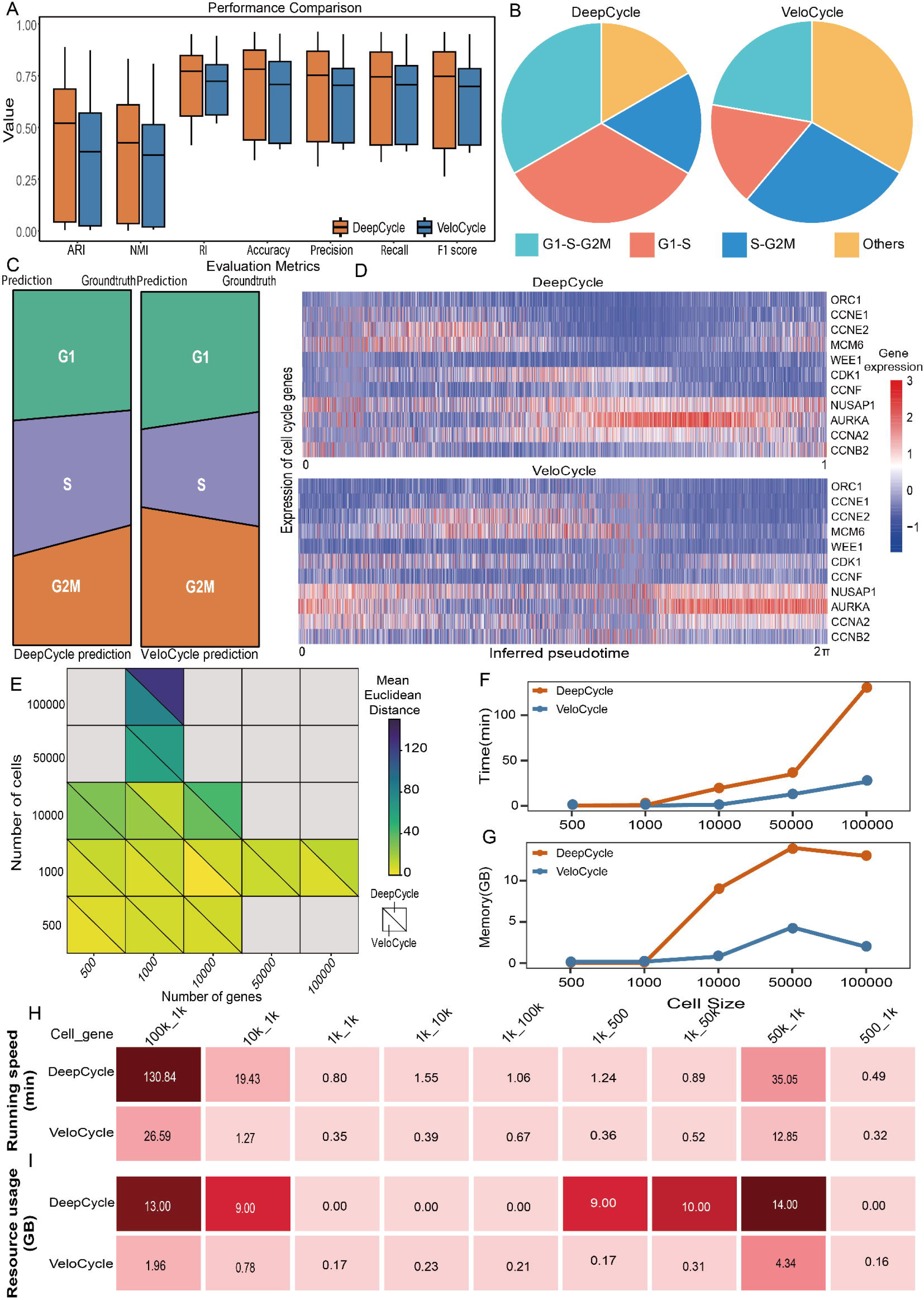
Comparison of cell-cycle velocity (and phase progression) inference by DeepCycle and VeloCycle. **A**, Boxplots summarize clustering- and classification-based metrics evaluated against reference phase labels comparing DeepCycle and VeloCycle across datasets. **B,** Pie charts show the inferred distribution of cell-cycle phase categories (G1-S-G2M, G1-S, S-G2M, and Others) for each method. **C,** Agreement with ground truth phase structure. Sankey plots compare the predicted phase partitioning (left: DeepCycle; right: VeloCycle) to the ground truth. **D,** Heatmaps display representative cell-cycle gene expression patterns ordered by inferred pseudotime. **E,** Heatmap summarizes the mean Euclidean distance between inferred pseudotime orderings under varying numbers of cells and genes. **F-G,** Runtime (**F**) and peak memory usage (**G**) as a function of cell number. **H-I,** Resource profiles across cell-gene configurations. Heatmaps report running time (**H**, minutes) and resource usage (**I**, GB) for a panel of dataset size combinations.

Both DeepCycle and VeloCycle correctly recover the canonical cell-cycle progression from G1 to S to G2M. However, phase-composition pie charts reveal clear differences in how accurately the two methods assign specific transition categories (**Fig. 4B**). When stratifying predictions into G1-S-G2M, G1-S, S-G2M, and other non-canonical transitions, DeepCycle assigns a substantially larger fraction of cells to the correct cell-cycle transition categories, indicating more precise identification of phase progression. In contrast, VeloCycle exhibits a markedly higher proportion of cells classified as “Others,” reflecting a higher rate of ambiguous or incorrect transition inference and overall weaker resolution of cell-cycle dynamics (**Fig. 4C**, **Supplementary Fig 2**).

The heatmaps show that both inferred pseudotime orderings align with canonical cell-cycle gene programs (**Fig. 4D**). Early-to-mid pseudotime is enriched for G1/S-associated genes such as ORC1, CCNE1, CCNE2, MCM6, whereas late pseudotime exhibits higher expression of G2/M-associated genes such as NUSAP1, AURKA, CCNA2, and CCNB2. Notably, VeloCycle’s pseudotime is represented on a circular coordinate from 0 to 2π, which matches the intrinsic periodicity of the cell cycle and visually reinforces closure of the trajectory. DeepCycle uses a linear 0-1 pseudotime scale that still captures the phase progression but expresses the cycle as a linear ordering rather than an explicitly closed loop.

We further evaluated velocity accuracy across different sizes of genes and cells using the Euclidean distance between inferred and ground-truth velocities. The mean Euclidean distance matrix indicates that errors remain low for both methods in small to moderate settings (**Fig. 4E**), but DeepCycle’s distances increase more visibly as cell numbers grow. This pattern suggests that DeepCycle may become more sensitive to optimization or resource constraints at larger scales, which can manifest as decreased pseudotime fidelity.

Scalability differences are pronounced and widen with dataset size. For 100k cells and 1,000 genes dataset, DeepCycle requires 130.84 minutes versus 26.59 minutes for VeloCycle (**Fig. 4F&H**), and 13.00 GB versus 1.96 GB of memory, respectively (**Fig. 4G&I**). For 10k cells and 1,000 genes, DeepCycle takes 19.43 minutes compared to 1.27 minutes for VeloCycle (**Fig. 4 F&H**). Across the reported configurations, DeepCycle shows a clear memory “jump” at moderate-to-large sizes, whereas VeloCycle’s memory footprint remains low and increases more smoothly with scale (**Fig. 4F&G**).

### Multi-omics velocity

#### Integrated epigenome-transcriptome velocity modeling

scKINETICS infers gene-specific kinetics by incorporating differential accessibility peaks ^7^, while MultiVelo is a paired multiome model that explicitly couples chromatin accessibility and RNA measurements from the same cells to estimate a joint temporal signal and infer state transitions ^6^. Across the embryonic mouse multi-omics dataset (**Fig. 5A**), both scKINETICS and MultiVelo recovered a broadly coherent developmental flow on the joint embedding, but scKINETICS showed a modest and more consistently favorable signal across complementary validity metrics (**Fig. 5B-D**). In particular, scKINETICS achieved higher directionality consistency as reflected by the distribution shift in CBDir, whereas MultiVelo displayed a wider spread and a more downward-shifted CBDir distribution, indicating more frequent local disagreements with the expected transition direction (**Fig. 5B**). Consistently, scKINETICS exhibited near-saturated within-neighborhood coherence in ICCoh and a consistency-score distribution concentrated toward higher values, while MultiVelo showed visibly greater dispersion, suggesting that its inferred velocity field is more sensitive to local variation in this dataset (**Fig. 5B**). By contrast, clustering metrics (inter-class distance, intra-class distance, and the inter-intra ratio) were largely comparable between the two methods, implying that both methods preserve a similar degree of between-state separation relative to within-state compactness, and that the observed advantage of scKINETICS primarily manifests in directionality and local coherence rather than global cluster geometry (**Fig. 5C&D**).

**Fig 5.**
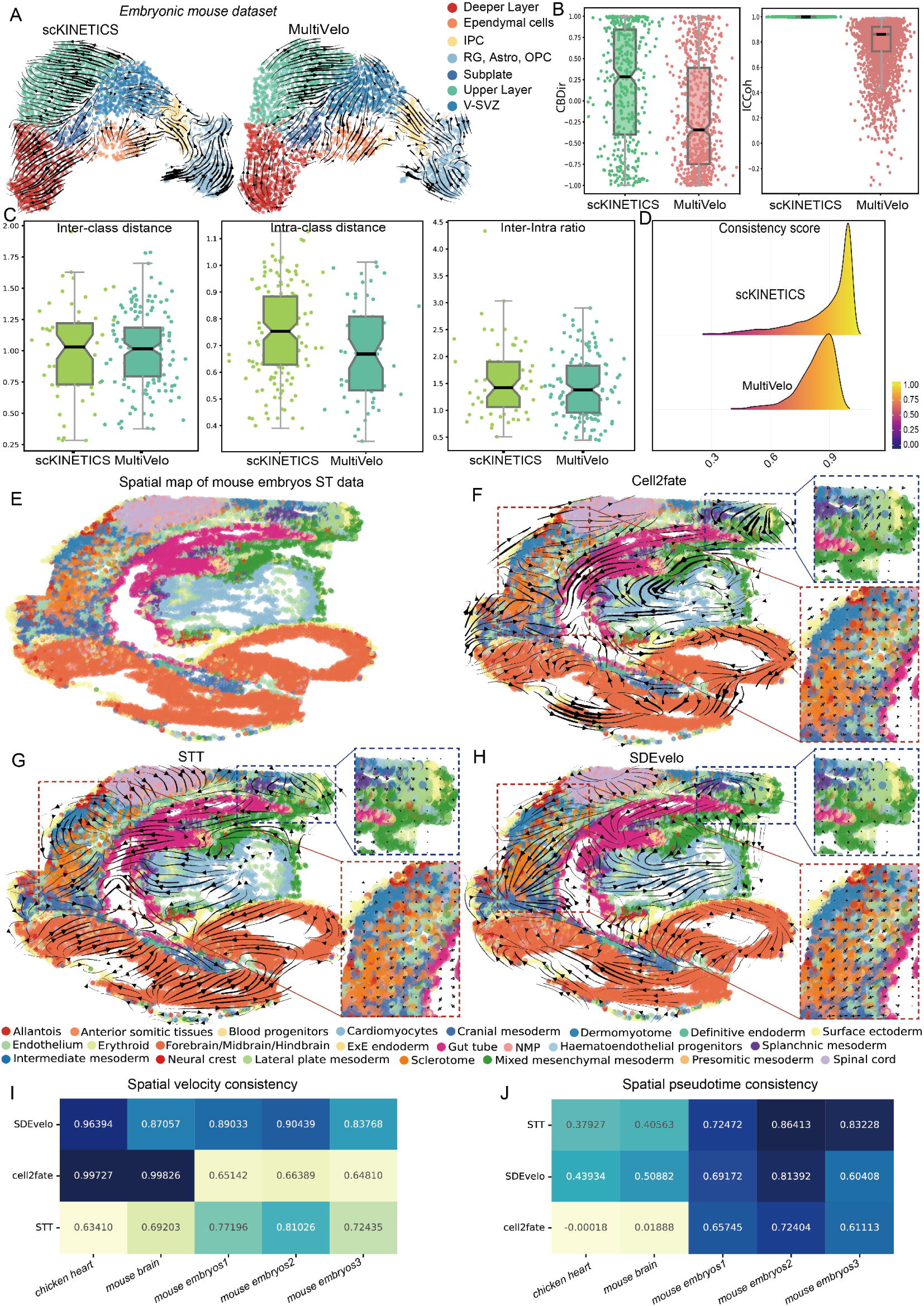
Benchmarking multi-omics velocity on various datasets. **A**, Streamline visualizations of the inferred flow on the joint embedding for scKINETICS and MultiVelo, with cells colored by annotated populations. **B,** Cross-boundary directionality (CBDir) and within-cluster coherence (ICCoh) comparing the two methods. **C,** Cluster-based metrics summarizing between-state separation and within-state compactness for each method. **D,** Distribution of the consistency score, reflecting local neighborhood smoothness of the velocity field. **E,** Spatial map of the mouse embryo ST dataset colored by annotated cell types. **F-H,** Spatial velocity streamlines overlaid on the same embedding for cell2fate (**F**), STT (**G**), and SDEvelo (**H**); dashed boxes highlight representative regions used for qualitative comparison from biological relevance. **I,** Spatial velocity consistency across multiple spatial datasets. **J,** Spatial pseudotime consistency across the same datasets.

Methodologically, these two approaches differ primarily in their data requirements and how epigenomic information is incorporated. scKINETICS can augment RNA-based velocity inference with regulatory priors derived from chromatin accessibility, such as peaks and motif information obtained from bulk or single-cell ATAC profiles, and therefore does not require paired RNA-ATAC measurements. In contrast, MultiVelo is designed for single-cell multi-omics datasets with matched RNA and chromatin profiles per cell and explicitly models the coupled epigenome-transcriptome dynamics, which constrains its applicability to paired multi-omics data. Accordingly, scKINETICS has broader practical coverage across datasets where only RNA is available together with external or aggregate chromatin-derived regulatory information. Finally, although Chromatin Velocity was initially considered and was showed in primary results, we did not include it in this general benchmarking framework because, in practice, it functions primarily as a specialized scVelo-style workflow tailored to scGET-seq data, rather than as a standalone multiome velocity method with outputs that are directly comparable to those of the broader set of evaluated tools.

#### Protein velocity inference method

Currently, only *protaccel* provides an explicit implementation of protein velocity. Rather than competing with RNA velocity estimators, *protaccel* proposes a complementary concept that uses simultaneously measured protein abundances to infer short-term changes at the protein layer, thereby offering a translational counterpart to RNA velocity. Conceptually, the method extends the RNA-velocity framework by coupling unspliced and spliced transcript counts with protein measurements to extrapolate cell states forward in time, and it further introduces protein acceleration to capture transient dynamics that may not be apparent from first-order changes alone. *protaccel* also supports joint visualization of RNA velocity and protein velocity in a combined stream plot (**Supplementary Fig. 3A**), which enables a qualitative assessment of whether transcriptional and proteomic dynamics are aligned or exhibit systematic divergence in specific regions of state space. Nevertheless, this framework remains a valuable and still underexplored direction. It directly addresses the well-recognized gap between transcript abundance changes and downstream functional effectors at the protein level, and it motivates future method development and evaluation criteria for multiomics velocity that go beyond RNA-only dynamics.

#### Spatial velocity inference methods

Using the mouse embryo spatial transcriptomics dataset, we first visualized the annotated tissue map on spatial coordinates (**Fig. 5E**, **Supplementary Fig. 3B**). We then inferred spatial velocity fields and overlaid streamlines for three spatial methods, cell2fate, STT, and SDEvelo (**Fig. 5F-H**, **Supplementary Fig. 3B**). Both STT and SDEvelo showed smooth and locally coherent flows that followed major tissue domains and changed gradually across boundaries, consistent with structured spatial progression at the tissue scale. In contrast, cell2fate produced several streamline segments that crossed sharp germ-layer boundaries in the zoomed region (e.g., trajectories pointing from mesodermal lateral plate areas toward surface ectoderm areas ^37,38^), which is failed to represent true fate transitions and suggests confounding from local spatial proximity and mixed signals rather than lineage progression (**Fig 5F**, blue box). Using the somitogenesis filter, SDEvelo also can recover a coherent PSM-anterior somitic tissues-dermomyotome/sclerotome progression compared to cell2fate and STT (**Fig 5F-H**, red boxes). In the corrected streamline field, vectors originate in the PSM domain, flow toward anterior somite regions, and then extend into dermomyotome and sclerotome areas ^39^. This matches the expected spatial ordering of somite maturation, indicating that SDEvelo can precisely model this differentiation path.

We next assessed cross-dataset robustness using spatial velocity consistency and spatial pseudotime consistency (**Fig. 5I-J**). SDEvelo achieved the highest and most stable spatial velocity consistency across datasets, while STT showed the strongest spatial pseudotime consistency on the three mouse embryo replicates, supporting its ability to recover reproducible spatial progression patterns in embryonic development. By comparison, cell2fate displayed marked dataset dependence, with substantially lower consistency on the mouse embryo replicates than on the other platforms (**Fig. 5 I-J**). These results indicate that, in embryonic spatial data, STT and SDEvelo better preserve biologically plausible, spatially coherent dynamics, whereas cell2fate may yield cross-boundary directions that should be interpreted cautiously.

### Challenges and practical guidelines

Based on the results of our systematic and comprehensive benchmarking, we summarize key limitations revealed by our results and outline practical guidelines (**Fig. 6**).

**Fig 6.**
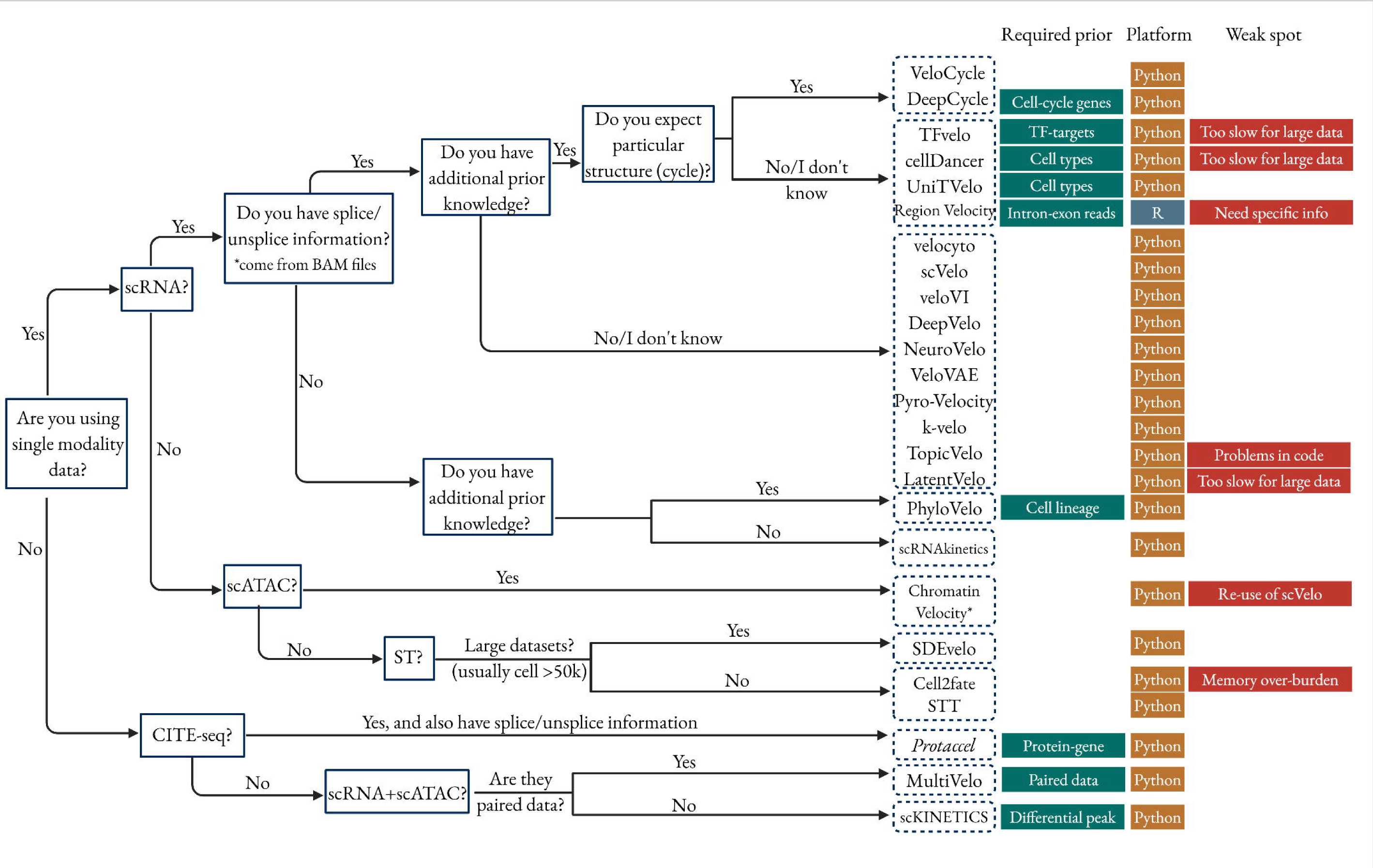
Guidance of the practical use for velocity inference tools.

#### Scalability remains a primary barrier for large datasets

A major bottleneck is non-completion at large problem sizes. In practice, many methods fail to return results once cell counts and gene counts grow beyond moderate regimes (e.g. larger than 10k cells), often due to memory pressure or long runtimes. Several methods cannot be run beyond moderate dataset sizes because they do not complete under the same time and memory budget. To make comparisons actionable, benchmarks should report completion and the largest tractable dataset size for each method, alongside runtime and memory under a fixed resource setting.

#### Gene selection can strongly modulate inferred velocity and inflate cell-to-cell variability

Velocity inference is highly sensitive to which genes are included in fitting and downstream vector field construction ^40^. Aggressive filtering, inconsistent gene panels across datasets, or reliance on highly variable genes can amplify apparent heterogeneity between cells, even when biological dynamics are smooth ^41,42^. This is particularly important because velocity is a direction vector defined in gene space: changing the gene basis changes the geometry of the vector field. A practical implication is that reproducible velocity analyses likely require standardized gene-selection protocols, sensitivity checks (e.g., stability under alternative gene sets), and reporting of gene-selection criteria as part of the main results rather than supplementary details.

#### RNA velocity can be systematically biased during fate commitment and intermediate states

A recurring biological limitation is that the standard interpretation of unspliced abundance as a proxy for impending upregulation can break down in intermediate or commitment states ^14,43^. In some transitional regimes, substantial unspliced mRNA may accumulate due to delayed processing, transcriptional bursting, altered splicing kinetics, or regulation that decouples transcription from maturation ^44–46^. This can bias inferred velocities toward “earlier” states, occasionally producing directions that contradict known biology. The implication is not that velocity is unusable, but that its assumptions are most fragile precisely at the states users often care about most: branching, commitment, and regulatory switching ^5,40^. Methodologically, this strengthens the case for models that incorporate uncertainty, condition-dependent kinetics, and explicit checks for assumption violations.

#### RNA velocity is often more reliable for trajectory visualization than for definitive fate determination

Streamline plots provide an intuitive entry point to RNA velocity, but they are ultimately a low-dimensional visualization of a high-dimensional vector field and can therefore mislead interpretation ^5,47^. The displayed flow depends on the embedding, neighborhood graph construction, smoothing, and plotting parameters, such that visually plausible streamlines may still deviate from the underlying direction in expression space, particularly when embeddings distort geometry or when local neighborhoods mix states ^40,48^. These projection and smoothing effects imply that RNA velocity, in its current form, is more reliable as an exploratory tool for visualizing plausible directions of change and generating hypotheses than as a standalone basis for definitive fate assignment ^40^. Accordingly, streamline visualizations should be paired with quantitative directionality checks, such as cross-boundary metrics or lineage-consistent angle statistics, and future tools should communicate uncertainty and projection artifacts explicitly rather than presenting a single deterministic flow. When fate-level claims are required, additional evidence, such as perturbation experiments, lineage tracing, time-resolved sampling, or orthogonal modalities, remains essential.

Our systematic and comprehensive benchmarking indicates that method choice should be guided first by data modality and biological priors, and then consider computation and robustness. For standard scRNA-seq with spliced/unspliced counts, start from general RNA velocity tools with higher accuracy, such as veloVI. However, for large cohorts prioritize methods that reliably complete within resource budget rather than those that only perform well on small datasets, such as veloVI, scVelo, and VeloVAE. For spatial data, resolution is decisive, spatial velocity is most helpful when measurements approximate single-cell states, whereas for spot-based Visium a strong RNA-only estimate followed by spatial mapping is often more reliable unless the spatial method models spot mixing. For multi-omics, use modality-specific tools, such as MultiVelo for paired RNA and ATAC, scKINETICS when pairing is unavailable, and *protaccel* for CITE-seq. Finally, treat streamlines as visualization rather than fate proof and pair them with quantitative directionality checks.

## Discussion

This study provides a systematic benchmarking of RNA velocity inference pipelines across accuracy, scalability, stability, and usability, clarifying that “good performance” is inherently multi-dimensional. Across both simulated and real datasets, we observed substantial method-to-method variabilities, reinforcing that velocity estimates depend strongly on modeling assumptions.

Given the results and the limitations, a key opportunity is multimodal modeling that treats additional modalities as part of the dynamical system. Protein-aware, chromatin-aware, or spatially regularized dynamics can, in principle, reduce ambiguity in intermediate states and connect velocity to downstream effectors ^5^. However, most current “multimodal” workflows remain RNA-first and mapping-second ^9^. Progress will require models that use multimodal measurements to constrain kinetics and directionality during inference, along with evaluation criteria that go beyond RNA-only agreement metrics.

We also found that spatial velocity performance depended strongly on the spatial platform and effective resolution (**Supplementary Fig. 3C**). In particular, spatially informed velocity models tended to yield clearer benefits on seqFISH datasets, whereas on spot-based 10x Visium data they provided worse results compared to RNA-only velocity tools with spatial mapping such as veloVI. This platform dependence points to a key methodological sensitivity that spatial velocity approaches may work better when spatial measurements approximate single-cell states and preserve sharp local boundaries, but they degrade when each spatial unit is a mixture of multiple cell states. SeqFISH typically provides single-cell or near single-cell resolution with higher cell-type purity and a more faithful neighborhood structure for graph-based spatial regularization ^49,50^. By contrast, 10X Visium spots aggregate multiple cells, so spot-level expression is inherently compositional ^51^. Under this regime, spatial smoothing can blur genuine transitions and can perform worse than an RNA-only estimator that is subsequently projected into space. These observations motivate future spatial-velocity development to incorporate mixture-aware modeling for spot-based technologies, to quantify spot purity and neighborhood ambiguity as part of routine quality control, and to avoid assuming that spatial adjacency necessarily implies state adjacency when spatial resolution is limited.

A practical outcome of this work is that velocity methods should be recommended conditionally. Different biological questions privilege different properties: boundary directionality for lineage progression, local coherence for stable vector fields, robustness for comparative studies, and efficiency for atlas-scale datasets. Future benchmarks of method developement should therefore define “trust regions” that specify when assumptions are plausible (e.g., strong splicing signal, limited mixing, adequate gene coverage, time-resolved sampling) and when velocity outputs should be treated as exploratory only.

Overall, our results support a cautious but constructive view that RNA velocity is a powerful descriptive tool for organizing dynamic hypotheses, yet its inferential claims remain fragile under large-scale computation, gene-selection sensitivity, projection artifacts, and assumption violations during fate commitment. The most promising path forward is not incremental tuning of visualization, but deeper modeling that integrates multimodal constraints and platform-specific measurement structure, especially for spatial transcriptomics where resolution and mixing fundamentally alter what “directionality” can mean.

## Methods

### Velocity inference methods

We gathered a list of 36 velocity inference methods by searching the literature for “RNA velocity” or “velocity” and “single cell” or “spatial”. Methods were excluded from the evaluation based on several criteria: 1) not completely and freely available for the code; 2) do not created or optimize the existed methods; 3) do not used for velocity inference; 4) unresolved errors during downloading and wrapping; We excluded ten tools from the benchmarking: CellPath ^52^, Cytopath ^53^, RNA-ODE ^54^, OTvelo ^55^, SIRV ^56^, scRegulocity ^57^, VeloSim ^58^, WF-Velo ^59^, VeTra ^60^, and VeloViz ^48^. Of these, CellPath, Cytopath, and RNA-ODE specialize in inferring trajectories using previous inferred RNA velocity information. OTvelo identified ‘gene velocity’ by using the change rate in different time points rather than using spliced/unspliced information for each cell. VeloSim was employed to generate simulation data for this study, while VeloViz is designed for visualizing RNA velocity. The detailed information of excluded methods can be found in Supplementary Table S2. We finally included 26 methods for the evaluation, including 20 methods for RNA velocity, 7 methods for multi-omics velocity (chromatin velocity, protein velocity, and spatial velocity), and 2 methods for velocity-based cell cycle inference (Supplementary Table S1).

### Datasets

#### Real datasets

We assembled 62 real scRNA-seq and spatial datasets via two complementary routes. (i) Tool-associated corpora. We collected the author-curated datasets shipped with or linked from widely used RNA-velocity toolkits (e.g., tutorial/benchmark sets in their repositories), ensuring that spliced/unspliced layers or raw reads were available. (ii) Repository search. We performed a systematic search of public archives (primarily GEO/ArrayExpress/SRA) using queries such as “single cell,” “single cell RNA/ATAC,” then selected studies whose original publications reported RNA velocity analyses. For each candidate accession, we confirmed (a) droplet- or plate-based scRNA-seq with recoverable spliced/unspliced counts or raw sequences; (b) the study captured a dynamic biological process amenable to velocity analysis, such as cell differentiation, organ development, and tumor progression. (c) each dataset contains all the information required by the evaluated tools. Spatial datasets were included when the authors provided velocity-ready matrices or when raw reads allowed construction of spliced/unspliced layers.

#### Synthetic datasets

We generated 114 synthetic single-cell datasets with known ground-truth velocities using Dyngen ^61^ (Supplementary Table S4). For each experiment, we instantiated both canonical developmental backbones and various backbones, including linear, bifurcating, bifurcating loop, consecutive bifurcating, trifurcating, cyclic and disconnect structure, sampled a gene-regulatory network and kinetic parameters (transcription, splicing, degradation) from Dyngen’s defaults, and simulated transcript dynamics via a stochastic reaction framework. We varied dataset size (ranging from 500 to 500,000 cells), the number of genes (ranging from 500 to 200,000 genes) and produced multiple random replicates per setting using different seeds (Supplementary Methods).

### Preprocessing steps

We preprocessed raw data according to assay type while preserving spliced/unspliced layers for velocity. For 10x Genomics scRNA-seq we ran CellRanger (https://github.com/10XGenomics/cellranger) by using the Grh38 and mm10 as reference genome to obtain UMI matrices and aligned BAM files. For plate-based protocols (e.g., Smart-seq2) we aligned with STAR and quantified exon/intron reads to derive spliced/unspliced counts ^62^. For Drop-seq/10x inDrop data we used dropEst ^63^. Then we used velocyto to obtain the splice/unsplice counts ^14^. Quality control removed low-quality cells using adaptive thresholds on detected genes, total UMIs, mitochondrial and ribosomal RNA fractions, and extreme unspliced fractions. We used the markers reported by original literatures to perform the cell type annotation. The processed datasets are h5ad or loom format. For highly variable gene (HVG) selection, we did not impose a unified external set; instead, each RNA velocity method used its own built-in gene selection routine with default parameters, and methods that do not require HVG selection were run on all expressed genes passing minimal detection thresholds. Detailed information for each method is shown in **Supplementary Methods**.

### Benchmarking metrics

We used multiple quantitative evaluation metrics to benchmark RNA velocity-based methods on real and simulated datasets.

### For RNA velocity inference tools

#### Consistency score

The consistency score ^64^ of cell *i* is defined as the mean correlation between its velocity vector *v_i_* and the velocity vectors of its *n* nearest neighboring cells, as follows:

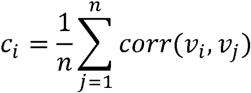

where *j* denotes a neighboring cell of cell *i*, and *n* is the total number of neighbors, with a default value of *n*=20. In our benchmarking, we used cosine similarity as the measure of correlation.

To evaluate whether the consistency scores are concentrated around a specific value, broadly dispersed, or exhibit multiple peaks, we fit a Gaussian distribution to the consistency scores across all cells. The peak of the distribution represents the most frequent value, indicating where most cells lie in terms of consistency. A higher peak suggests that a larger number of cells have velocities that are more consistent with those of their neighbors, reflecting better performance. Therefore, we use the peak value of the fitted distribution to rank the compared methods: a higher peak corresponds to a higher ranking.

#### Velocity Angle consistency

Angle consistency measures the alignment between the predicted velocity direction and the expected direction, defined by the angular difference between the two. Specifically, the included angle is the angle between the observed velocity vector and the expected reference direction.

To define the expected direction, we computed the centroid of each cell type and constructed the lineage by several biology experts, guided by known biological trajectories to serve as the ground truth. For each cell, we calculated both its observed velocity direction and its expected direction, using cosine distance to quantify their angular separation.

Angle consistency is visualized using rose plots, which display the distribution of cosine distances (i.e., angular deviations) between each cell’s velocity vector and the reference direction computed from pseudotime ^65^. Each cluster is plotted separately with 20 bins. Angles close to 0° (i.e., < 90°) indicate that the velocity is aligned with the intended direction, while angles near 180° indicate movement opposite to the expected trajectory.

To quantitatively compare methods, we computed the proportion of cells with angles between 0° and 60°. A higher proportion indicates better alignment and therefore better performance, which is used to rank the methods accordingly.

(for each cell l, we choose its neighbor m which is the furthest along in pseudotime (i.e. generally most duct-like) and then compute the direction spanned from cell l to that neighbor.)

#### Inter-class distance

The inter-cluster distance measures how far apart different clusters are. It reflects the separation between cluster centers. Larger values indicate better separation. It is defined as:

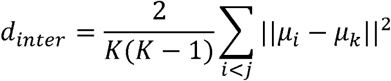

where *K* is the total number of clusters, and μ_*i*_ and μ_*j*_ are the centroids of clusters *C_i_* and *C_j_*, respectively. The condition *i* < *j* ensures that each pair of clusters is counted only once.

#### Intra-class distance

The intra-cluster distance measures how close the cells within the same cluster are to each other. It reflects the compactness of each cluster. Smaller values indicate better compactness. It is defined as:

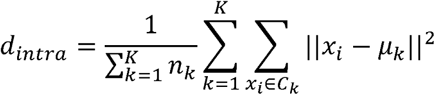

where *K* is total number of clusters, *n_k_* is number of cells in cluster *C_k_*, μ_*k*_ is the centroid of cluster *C_k_*, and *x_i_* is a cell belonging to cluster *C_k_*.

#### Inter/Intra ratio

The inter/intra ratio quantifies the overall clustering quality by comparing between-cluster separation to within-cluster compactness. Larger values indicate better clustering performance. It is defined as:

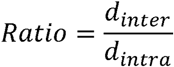

where *d_inter_* is the average inter-cluster distance, and *d_intra_* is the average intra-cluster distance, as defined above.

#### Cross-boundary direction correctness (CBDir)

CBDir ^66,67^ evaluates the correctness of predicted transitions from a source cluster to a target cluster by focusing on boundary cells, using ground truth directionality. The boundary of a source cluster refers to those cells whose neighbors include cells from the target cluster, and vice versa. Boundary cells are chosen because they represent transitional states that reflect biological development over a short time period. CBDir is computed as:

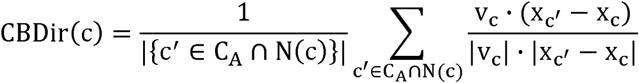

where C_A_ *is the set of cells in the target cluster* A, N(c) denotes the neighbors of cell c, v_c_ is the RNA velocity vector of cell c. x_c_ *and* x_c′_ are the low-dimensional coordinates of cells c *and* c′. The vector x_c′_ - x_c_ represents the expected short-term displacement in the latent space. This metric requires known ground truth transitions to define the source and target clusters and is used to assess how well the predicted velocities align with biologically meaningful transitions, thereby reflecting the reliability of the RNA velocity model. Higher values indicate better performance.

#### In-cluster coherence (ICCoh)

ICCoh ^66,67^ is calculated as the average cosine similarity of RNA velocity vectors among cells within a homogeneous cluster. It reflects the smoothness and consistency of velocity directions inside the cluster. Note that ICCoh can yield a high score even if some velocity directions are reversed, as it measures similarity regardless of absolute orientation. Formally, it is defined as:

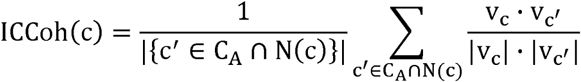

where v_c′_ is the RNA velocity vector of cell c′.

#### Ground-truth correlation

For simulated datasets with known ground-truth velocity, we quantified method performance by comparing the inferred velocity to the ground truth using the cosine similarity. For each cell t, we treated the inferred and ground-truth RNA velocity as vectors in gene-expression space and computed

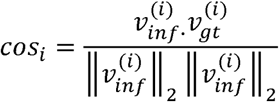

where 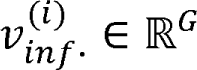 is the inferred velocity vector for cell *i* across the *G* genes, and 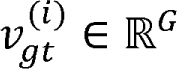 is the corresponding ground-truth velocity vector. We then mapped the cosine similarity to [0,1] as

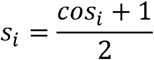

This metric captures directional agreement between inferred and true velocities while remaining insensitive to global scaling differences. A score of *s_i_* = 1 indicates perfect alignment, and larger values indicate better agreement.

### For velocity inference and cell-cycle prediction tools

For the velocity-based cell-cycle prediction tools, we specifically evaluated DeepCycle ^35^ and VeloCycle ^36^ on real and simulated datasets with cell-cycle labels. For each dataset, the predicted cell-cycle labels (G1, S, G2/M) from the two methods were first compared with the ground-truth labels using a panel of metrics, including adjusted Rand index (ARI), normalized mutual information (NMI), Rand index (RI), accuracy, precision, recall, and F-score. In all these metrics, higher values indicate better agreement between predicted and true cell-cycle states. In addition to these discrete metrics, we also took advantage of the continuous cell-cycle position estimated by both methods. For each cell, we represented the true cycle position and the predicted position as points on the unit circle and computed their Euclidean distance. Moreover, to assess the biological plausibility of the predictions, we examined the expression of canonical cell-cycle-related genes under the states inferred by each method. Using curated G1, S and G2/M gene sets, we computed average expression for each gene set within the predicted phases and along the predicted continuous cycle. We then assessed whether G1, S and G2/M markers peaked in the expected regions of the cycle and whether their expression trends were better aligned with the ground-truth cycle under DeepCycle or VeloCycle. Together, these complementary criteria, discrete agreement with true labels, Euclidean distance to the true cycle position, and phase-specific expression of cell-cycle genes were used to compare the overall performance of DeepCycle and VeloCycle.

### For multi-omics velocity inference tools

Current multi-omics velocity inference methods are highly heterogeneous in both modeling assumptions and output formats, and their results are not directly comparable within a single unified framework. Therefore, in this section we focus on how adding non-transcriptomic information affects velocity estimation. Specifically, we considered three classes of multi-omics velocity approaches: (i) scRNA + scATAC / ATAC methods that couple RNA kinetics with chromatin accessibility; (ii) scRNA + protein (CITE-seq) methods that augment RNA dynamics with protein acceleration; and (iii) scRNA + spatial transcriptomics methods that integrate local spatial context into velocity estimation. For each class, representative tools were run on matched multi-omics datasets using the authors’ recommended preprocessing and default parameters. We then compared multi-omics velocities to these two references across multiple datasets, examining the changes in the reconstructed trajectories and velocity stream after adding ATAC, spatial, or protein information, and the representation format of multi-omics velocity such as per-gene kinetic parameters, and cell-wise phase/latent time, to assess both the numerical impact and the interpretability of multi-omics velocity inference.

For spatial velocity inference tools, we construct two measurements to evaluate the accuracy:

### Spatial Velocity Consistency

Spatial velocity consistency ^68^ measures the local agreement of RNA velocity vectors in space. Given precomputed RNA velocities *V* = {*v*_1_, …, *v_n_*} and spatial coordinates, it computes the Pearson correlation of velocity vectors among each cell’s *k* nearest neighbors, quantifying how consistent velocities are across the tissue spatial neighborhood:

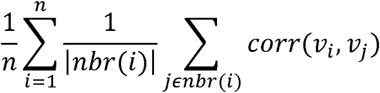

### Spatial Time Consistency

Spatial time consistency ^68^ quantifies the spatial coherence of inferred cell times. Given a spatial graph with adjacency matrix *A*ϵ{0,1}^*n×n*^ and pseudo times *T* = {*t*_l_, …, *t_n_*}, it is defined as the Moran’s I statistic of cell times over the spatial coordinates, measuring the degree to which neighboring cells have similar temporal states.

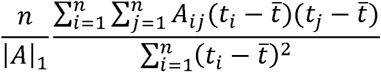

### Methods execution

Each execution of a method on a dataset was performed separately as a job. For the real datasets and the simulated datasets used for metric measurement, the tasks were allocated with independent R/Python sessions, or an error was produced, a zero score was returned for that execution and the error information will be recorded (**Supplementary Methods**). Specifically, for the tools that not fit for metrics calculation, the rank will be set as “out-of-rank”.

### Docker construction

All Docker images were built upon the official NVIDIA CUDA base image (nvidia/cuda: v11.8.0-devel-ubuntu v22.04) to ensure consistent GPU driver compatibility and stability. For each analytical method, a dedicated, self-contained Docker image was constructed by following the installation guidelines provided in its official documentation and source code repository. These images encapsulate all necessary libraries and dependencies. All custom-built images have been made publicly available on Docker Hub for accessibility and reproducibility.

### Hardware Deployment and Resource Allocation

The Docker environments were deployed on two high-performance servers with the following specifications: GPU-Accelerated Server: Equipped with dual Intel® Xeon® Silver 4210R CPUs (total 20 cores/40 threads, 2.40-3.20 GHz) and 128 GB of RAM. This server is augmented with two NVIDIA® GeForce® RTX 3090 GPUs, each providing 24 GB of GDDR6X VRAM for massively parallel computations and deep learning tools. CPU Server: Equipped with dual AMD EPYC™ 7C13 CPUs (total 128 cores/256 threads, 1.50-3.70 GHz) and 128 GB of RAM. This server is designated for computationally intensive tasks such as statistical-model-based tools and machine-learning-based tools.

Therefore, our deployment strategy allocated methods requiring CUDA acceleration to the GPU-Accelerated Server, while all other methods were run on the High-Core-Count CPU Server. To estimate the scalability and computational cost of tools, we enforced strict resource limits for each container: on the CPU Server, each container was allocated 32 GB of RAM with 32 CPU cores. On the GPU Server, each container was allocated 32 GB of RAM with 8 CPU cores. To prevent performance degradation from resource contention, all allocated cores were pinned to a single physical NUMA (Non-Uniform Memory Access) node each container. Furthermore, each container was granted exclusive access to a dedicated physical GPU to prevent resource sharing conflicts.

### Scalability

To assess the scalability of each method, we used Dyngen to generate a panel of synthetic RNA-velocity datasets with increasing size. We varied the number of cells and genes so that the datasets spanned several orders of magnitude (from hundreds to hundreds of thousands of cells and from ∼10³ to ∼10l genes) with multiple topological structures.

For every run, we monitored computational cost and resource usage. Specifically, we recorded the wall-clock runtime from the RNA velocity estimation processes, the method-specific preprocessing such as data format transfer, normalization, gene filtering, neighbor-graph construction was excluded. In parallel, we tracked peak CPU usage (%), peak resident memory (GB) and, for GPU-accelerated implementations, peak GPU memory (GB) using system-level profiling tools. Runs that crashed or exceeded the available hardware resources were marked as failures for that configuration and excluded from aggregate statistics. This approach can provide a clear assessment of each method’s computational efficiency.

### Stability

We assessed the stability of velocity methods along two axes: down-sampling robustness and multi-batch reproducibility.

### Down-sampling robustness

To evaluate sensitivity to data loss and noise, we used the bifurcating dataset with 1,000 cells and 10,000 genes (“bifurcating_cell1000_gene10000”) generated by Dyngen as a reference. From this dataset, we created down-sampled datasets by randomly retaining 60%, 70%, 80%, 90%, and 95% of cells or genes (sampling without replacement, separately for cells and genes). For each down-sampled dataset and each method, we reran the full velocity-inference pipeline using the same preprocessing and parameter settings as on the full dataset. Stability was quantified by comparing the inferred velocity on each down-sampled dataset to the ground-truth velocity using the cosine distance metric described above, computed on the intersection of cells and genes shared with the original dataset. For each down-sampling level, we summarized performance by the mean/median cosine distance across cells; methods that showed small degradation of this metric relative to the full data were considered more stable to subsampling.

### Multi-batch reproducibility

To assess the impact of stochastic optimization and random initialization, particularly for deep learning-based methods, we performed repeated runs on one real dataset (7_Mm_PancreaticE15.5) and one simulated dataset (bifurcating_cell1000_gene10000). For each deep learning method and each dataset, we executed the full pipeline five times with different random seeds. Let 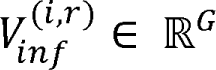 denote the velocity matrix (cells × genes) from run *r* ϵ {1,2,…5}. For every pair of runs (*r*_1_,*r*_2_), we first restricted to the set of cells present in both runs and then quantified run-to-run agreement by the mean cosine similarity of per-cell velocity vectors:

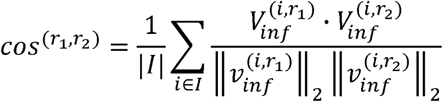

where *I* denotes the set of shared cells between runs *r*_1_ and *r*_2_. We then mapped *cos*^(*r*_1_,*r*_2_)^ to [0,1] as

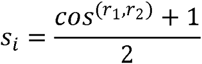

The multi-run reproducibility score for a method on a given dataset was defined as the mean of *s*^(*r*_1_,*r*_2_)^ over all 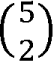 run pairs. Higher values indicate greater reproducibility across independent runs.

### Usability

To evaluate the usability of RNA-velocity tools, we combined computational experts review with an AI agent (GPT-4). For each method, we inspected the official installation instructions, documentation and tutorials, example notebooks or pipelines, issue tracker, and code repository, and scored the tool on five dimensions (1-10 scale, higher is better):

- **Install-friendly score**: ease of installation and dependency management, presence of packaged releases or containers, and clarity of setup instructions.
- **Usage guidance score**: quality and completeness of documentation, tutorials, example workflows, and in-line help.
- **Maintenance score**: apparent level of maintenance, including recency of commits, responsiveness to issues, and versioning practices.
- **Code quality score**: readability and structure of the codebase, modular design, presence of tests and type hints, and clarity of configuration.
- **Feedback score**: informativeness of error messages, logging, progress bars, and overall user feedback during a typical run.

The final scores reported in the usability table correspond to the average of ratings for each dimension combining the AI-agent and manual adjustments.

## Data and code availability

The datasets used for benchmarking were uploaded in Zenodo (https://zenodo.org/records/18102832). The code, docker files and the usage guideline are available in GitHub (https://github.com/vikkihuangkexin/VelocityBenchmarking) and our website (https://relab.xidian.edu.cn/RNAVelocity/).

## Acknowledgement

This work was partially supported by the National Institutes of Health [R01LM014156, R01GM153822, R01CA241930, R01AA032723-01 to X.Z], the National Science Foundation [2217515, 2326879 to X.Z], the Cancer Prevention and Research Institute of Texas [RP250043 to X.Z], and the Dementia Prevention and Research Institute of Texas [P018230 to J.L]. Dr. Liyu Huang is supported by the National Natural Science Foundation of China under grant Numbers 82227802 and 62373292. The funders had no role in study design, data collection and analysis, decision to publish or preparation of the manuscript.

---

**Supplementary Fig 1. Real-data accuracy ranking across proxy metrics.** Bar plots show the metric-specific rank of RNA velocity methods evaluated on real datasets. Higher values indicate better rank for the corresponding metric.

**Supplementary Fig 2. Cell-cycle phase inference across datasets.** Boxplots compare DeepCycle (top) and VeloCycle (bottom) predictions of cell-cycle phase scores across representative real datasets and cycle-simple synthetic datasets. Cells are grouped by ground-truth cell-cycle phase (x-axis), and the y-axis shows the corresponding method-predicted phase score.

**Supplementary Fig 3. Results of protein velocity and spatial velocity. A,** streamline showing velocity vectors estimated from RNA velocity, protein velocity, and their combined signal. Dots represent cells colored by broad immune cell types. **B,** spatial map of the 10X Visium chicken heart spatial transcriptomics dataset with spot annotations (left). Streamline visualizations of spatial velocity fields inferred by cell2fate, STT, and SDEvelo are overlaid on the same spatial coordinates (right and bottom). **C,** quantitative comparison between spatial velocity and RNA velocity tools across datasets. Left: spatial time consistency. Right: spatial velocity consistency.

