## Supplementary for "Benchmarking algorithms for RNA velocity inference": Supplementary Methods.docx

**Data generation**

**Mertric-used data generation**

For the main benchmarking metrics, we generated synthetic single-cell datasets using Dyngen under predefined trajectory topologies and matched combinations of cell and gene numbers. For each configuration, we sampled an independent random seed for every simulation run to avoid repeated realizations and to ensure that the inferred velocities were evaluated on non-identical stochastic outcomes. The resulting datasets provide internally consistent regulatory dynamics together with the associated ground-truth quantities required for accuracy and stability evaluation.

**Extreme-large data generation**

Dyngen does not natively support simulations beyond approximately 10,000 cells in our setting. To stress-test computational scalability at larger scales, we constructed three extra-large datasets by merging multiple dyngen-generated datasets of the same configuration to reach the target cell counts. Because these merged datasets do not preserve a single, rigorously simulated regulatory program or a coherent ground-truth velocity field across the full combined population, they were used exclusively for scalability analyses such as completion, runtime, and resource usage, and were not included in accuracy-based evaluations.

**Down-sampling data generation**

To evaluate robustness to data loss and noise, we performed controlled down-sampling on a representative simulated dataset with a bifurcating topology at 1,000 cells by 10,000 genes. We randomly subsampled both cells and genes to retain 60%, 70%, 80%, 90%, and 95% of the original data, and then re-ran each method using the same processing and evaluation workflow. These down-sampled datasets were used specifically for stability assessment under reduced coverage and increased stochasticity.

**Method execution**

**Velocyto**

As a pioneering approach, Velocyto established the framework for estimating single-cell RNA velocity by leveraging the ratio of spliced to unspliced RNA abundances. It employs a model based on first-order differential equations to fit transcription and degradation rates, thereby inferring future cellular states. However, its reliance on a strict steady-state assumption is often too idealized for complex biological dynamics. Furthermore, the tool is restricted to processing loom format files generated by its own pipeline, necessitating data conversion when working with other input formats.

**scVelo**

scVelo represents a comprehensive advancement in velocity inference, offering a stochastic model (which improves upon steady-state assumptions via second-order moments) and a dynamical model. The latter solves full transcriptional dynamics using maximum likelihood estimation to infer latent time and kinetic parameters without assuming steady states. It is highly compatible with the Scanpy ecosystem and diverse data formats.

**CellDancer**

CellDancer employs deep neural networks (DNNs) to infer non-linear velocity functions directly from neighboring cells, uniquely allowing for cell-specific reaction rates. While capable of modeling complex kinetics, it is computationally intensive, requiring GPU acceleration for efficient training. Additionally, it requires tabular data input, which can present data conversion challenges for high-dimensional datasets.

**VeloVI**

VeloVI introduces a probabilistic approach using deep generative models and variational inference. By estimating posterior distributions rather than point estimates, it provides uncertainty quantification for velocity vectors. It is GPU-dependent (PyTorch-based) and integrated into the scVI-tools library.

**UnitVelo**

UnitVelo is a spliced-RNA-oriented framework that utilizes Radial Basis Function (RBF) kernels and a global optimization term to resolve temporal inconsistencies between genes. We recommend using the 'unified-time' mode as the default setting to ensure synchronized inference. While GPU acceleration is highly recommended to significantly reduce computational time, processing datasets with large cell numbers may encounter GPU memory bottlenecks. In such instances, we advise utilizing the tool's downsampling function—specifically downsampling by cell type—to compute velocities on subsets and subsequently reconstruct the global velocity matrix.

**LatentVelo**

LatentVelo is a deep learning-based RNA velocity method that embeds cells into a low-dimensional latent space using a variational autoencoder. Differentiation dynamics are modeled in this latent space with neural ordinary differential equations, allowing inference of latent regulatory states that govern individual cell dynamics. This enables modeling of multiple lineages and prediction of latent trajectories, describing developmental paths of single cells rather than only local RNA velocity vectors. The dynamics-based latent embedding also performs batch correction, improving over standard autoencoder approaches that do not account for gene expression dynamics.

**Pyro-Velocity**

Pyro-Velocity is a probabilistic RNA velocity method that models multivariate gene expression to estimate the uncertainty of future cell states. It provides statistically grounded and interpretable information on cell fate and dynamics, enabling robust inference of RNA velocity in single-cell transcriptomics data.

**SDEvelo**

SDEvelo is a generative RNA velocity method that models the dynamics of unspliced and spliced RNAs using multivariate stochastic differential equations. It explicitly accounts for the inherent stochasticity of transcriptional dynamics while estimating a cell-specific latent time across genes. This enables robust inference of RNA velocity, accurate modeling of random dynamic patterns, and downstream analyses in both single-cell RNA-seq and spatial transcriptomics data.

**VeloAE**

VeloAE is an autoencoder-based RNA velocity method designed to denoise and robustly estimate cell transitions from single-cell transcriptomics data. It learns a low-dimensional representation of RNA velocity, improving stability and interpretability of inferred trajectories. VeloAE accurately captures stimulation dynamics and differentiation pathways across diverse biological systems, enhancing the utility of RNA velocity for studying transient cellular processes.

**PhyloVelo**

PhyloVelo is a computational RNA velocity framework that leverages monotonically expressed genes (MEGs) to reconstruct transcriptomic velocity fields. By combining with lineage information, it accurately infers linear, bifurcated, and convergent differentiation trajectories. PhyloVelo demonstrates high robustness across multiple lineage-traced datasets and identifies conserved gene functions associated with translation and ribosome biogenesis.

**STT**

STT is a spatial RNA velocity method that integrates mRNA splicing with spatial transcriptomics through a multiscale dynamical model. It reconstructs cell-state-specific dynamics and spatial transitions by learning a four-dimensional transition tensor and performing spatially constrained random walks. STT accurately captures multistability, short- and long-term transition paths, and associated genes across diverse biological systems and spatial scales.

**scRNAKinetics**

scRNAkinetics is a single-cell RNA kinetics inference method that estimates transcription, splicing, and degradation rates for each cell and gene using pseudo-time derived from various biological priors. It accurately models complex dynamic systems, including branching and time-dependent kinetics, and reveals coupling between transcription and splicing. The method is implemented with JAX for efficient optimization and scalable computation on large scRNA-seq datasets.

**κ-velocity**

κ-velocity improves RNA velocity estimation from single-cell RNA-seq by addressing biases and limitations in existing approaches. It introduces strategies for more accurate and robust inference of cell state velocities, enhancing the modeling of transcriptional dynamics and developmental trajectories.

**TopicVelo**

TopicVelo is a probabilistic RNA velocity method that uses a topic model to disentangle simultaneous, bursty transcriptional dynamics. It infers process-specific velocities by focusing on cells and genes associated with individual processes and integrates these signals into a global transition matrix. TopicVelo accurately captures complex cell-state transitions, transient dynamics, and terminal states, providing interpretable insights into multifaceted cellular functionality.

**Region Velocity**

Region Velocity is an RNA velocity method that estimates transcriptional dynamics at the sub-gene level by separately quantifying exonic and intronic expression. It models region-specific velocity using exon and intron count matrices, enabling finer resolution of transcriptional regulation. We followed the protocols.io tutorial (<https://www.protocols.io/view/region-velocity-estimation-and-visulization-with-s-eq2lyrejrvx9/v1>) and used the provided example dataset. Region velocity estimation was performed using the gene.region.velocity.estimates function with parameters theta.s = TRUE, fit.quantile = 0.05, and kCells = 10. Subsequent expectation-maximization (EM) refinement was conducted via gene.EM.velocity.estimates. The resulting velocity matrix and EM-inferred pseudotime were exported for downstream evaluation. Due to its requirement for separate exon and intron quantification, this method was applied to only one real dataset.

**VeloVAE**

VeloVAE is a variational autoencoder-based RNA velocity method that learns a low-dimensional latent representation of splicing dynamics. It models the joint distribution of spliced and unspliced counts through a probabilistic framework, enabling robust velocity inference and uncertainty quantification. We followed the example notebook on the VeloVAE GitHub repository (https://github.com/welch-lab/VeloVAE). For real datasets, preprocessing was performed using the velovae.preprocess function with the number of highly variable genes dynamically set according to data scale (200–3,000 genes) and the number of principal components set to 15 for small datasets (< 500 cells) or 30 otherwise. The standard VAE model was trained with dim_z = 5, tmax = 20, and default hyperparameters (full_vb = False, discrete = False). For simulated datasets, preprocessing thresholds were relaxed (min_shared_counts = 0, min_shared_cells = 0) to accommodate the characteristics of synthetic data. The latent dimension was reduced to dim_z = 4 given the smaller number of cell types in simulated data, with the number of highly variable genes (500–5,000) and principal components (30–50) adjusted according to data scale. Batch sizes were dynamically configured based on cell counts (64 for < 1,500 cells, 128 for < 15,000 cells, 256 otherwise). All other training parameters remained at their default values.

**DeepVelo**

We followed the guidelines on the DeepVelo GitHub repository: https://github.com/bowang-lab/DeepVelo/blob/main/README.md. We set the parameter n_var_genes=2000. DeepVelo is a graph convolution network-based RNA velocity framework that generalizes RNA velocity analysis to heterogeneous cell populations with time-dependent kinetics and multiple lineages, overcoming the limitations of predefined dynamics and cell-agnostic constant kinetic rates in traditional methods. It infers time-varying cellular rates of transcription, splicing and degradation, accurately recovers cell differentiation stages, and identifies functionally relevant driver genes regulating these dynamic processes. DeepVelo enables robust investigation of complex differentiation and lineage decision events in diverse developmental and pathogenic contexts using heterogeneous single-cell RNA-seq data.

**NeuroVelo**

We followed the guidelines on the NeuroVelo GitHub repository: <https://github.com/idriskb/NeuroVelo>. NeuroVelo is a computational method for reconstructing temporal cellular dynamics from static single-cell transcriptomics, which couples optimal linear projection learning with non-linear Neural Ordinary Differential Equations. It leverages dynamical systems theory to model time-dependent biological processes, enabling identification of genes and mechanisms driving temporal cellular dynamics. NeuroVelo exhibits high predictive power and outperforms state-of-the-art methods in mechanistic inference, while allowing direct inference of gene regulatory networks that govern cell fate from single-cell data.

**TFvelo**

We followed the guidelines on the TFvelo GitHub repository: https://github.com/xiaoyeye/TFvelo.

TFvelo is a computational RNA velocity framework that expands the RNA velocity concept by introducing gene regulatory information, without relying on splicing signals from unspliced/spliced mRNA. It leverages transcriptional regulation-driven dynamics to accurately fit gene dynamics in phase portraits and effectively infers cell pseudo-time and differentiation trajectories from RNA abundance data. TFvelo enables RNA velocity modeling across various single-cell datasets with improved robustness and accuracy compared to traditional splicing-dependent methods.

**cell2fate**

We followed the guidelines on the cell2fate website: https://cell2fate.readthedocs.io/en/latest/. We set the parameter n_var_genes=2000, and for the simulated data, we set min_shared_counts=0.

cell2fate is a Bayesian RNA velocity framework formulated based on the linearization of velocity ordinary differential equations (ODEs), enabling the solution of a biophysically more accurate model while avoiding coarse simplifications or numerical approximations of traditional methods. It decomposes RNA velocity solutions into modular components, establishing a biophysical link between RNA velocity inference and statistical dimensionality reduction. cell2fate enhances the interpretability of RNA velocity results, excels at reconstructing complex dynamics and capturing weak dynamical signals in rare or mature cell types, and facilitates the integration of spatial tissue architecture with transcriptional temporal dynamics in real-world single-cell studies.

**MultiVelo**

We followed the guidelines on the MultiVelo website: https://multivelo.readthedocs.io/en/latest/.

MultiVelo is a computational RNA velocity framework that extends classical RNA velocity by incorporating epigenomic data into a differential equation model of gene expression. It leverages a probabilistic latent variable model to estimate the switch time and rate parameters of chromatin accessibility and gene expression. MultiVelo improves the accuracy of cell fate prediction compared to RNA-only velocity methods and enables quantification of temporal relationships between epigenomic and transcriptomic dynamics.

***protaccel***

We followed the guidelines on the protaccel GitHub repository: <https://github.com/pachterlab/GSP_2019/tree/master>. *protaccel* is a computational RNA velocity framework that integrates simultaneous protein and RNA quantification from multimodal single-cell experiments. It leverages estimates of unprocessed transcript and protein abundances to extrapolate the past, present and future states of cells from static experimental snapshots. protaccel achieves consistent reconstruction of cell landscapes and phase portraits across multiple independent datasets and enables temporal dynamic analysis of multimodal single-cell data via a dedicated Python package.

**scKINETICS**

scKINETICS is an RNA velocity method that integrates single-cell RNA-seq with chromatin accessibility data to infer gene regulatory network (GRN)-constrained transcriptional kinetics. It leverages ATAC-seq–derived regulatory information to constrain velocity estimation, providing mechanistic insights into transcription factor–target relationships. We followed the demonstration notebook on the scKINETICS GitHub repository (https://github.com/dpeerlab/scKINETICS). Input scRNA-seq data were normalized to median library size, filtered to remove zero-expression genes, and log-transformed using scanpy.pp.log1p. Neighborhood graphs were computed with 15 neighbors and 40 principal components. For GRN construction, preprocessed differential accessibility peaks (width < 2,000 bp, obtained through differential analysis such as DESeq2) were annotated using the mm10 genome reference, and motif calling was performed with a p-value threshold of 1 × 10^-10. Target annotations were prepared for each cell cluster with MAGIC imputation for data smoothing. The expectation-maximization algorithm was executed with 15 parallel threads and a maximum of 20 iterations. Velocity graphs were constructed using 30 nearest neighbors for embedding. Due to its requirement for preprocessed differential accessibility peaks from ATAC-seq, this method was applied to only one real dataset.

**VeloCycle**

We followed the guidelines on the VeloCycle GitHub repository: https://github.com/lamanno-epfl/velocycle/tree/main/tutorials.We set the parameter num_steps=1000.

VeloCycle is a Bayesian RNA velocity framework that couples velocity field and manifold estimation in a unified, reformulated framework to identify parameters of an explicit dynamical system. It specializes in modeling gene regulation dynamics on one-dimensional periodic manifolds, particularly the cell cycle, and achieves statistically consistent inference with improved robustness over heuristic-based RNA velocity methods. VeloCycle enables accurate estimation of cell cycle periods and reveals dynamic differences in progenitors or gene perturbation contexts via a modular single-cell RNA-seq analysis toolkit.

**DeepCycle**

We followed the guidelines on the DeepCycle GitHub: https://github.com/andreariba/DeepCycle. We set the parameter expression_threshold=0.5. DeepCycle is a deep learning-based approach for single-cell RNA-seq data analysis, specifically designed to fit the cycling patterns of cell cycle-related genes in the unspliced-spliced RNA space. It enables the construction of high-resolution transcriptomic maps of the entire cell cycle and facilitates systematic characterization of cell cycle stages by capturing gene regulation dynamics without external cellular perturbation. DeepCycle exhibits efficient measurement of cycling gene dynamics and is applicable to multiple cellular models for cell cycle studies in diverse biological contexts.
