## Supplementary figures and images for "Benchmarking algorithms for RNA velocity inference"

### Supplementary Figure 1.png

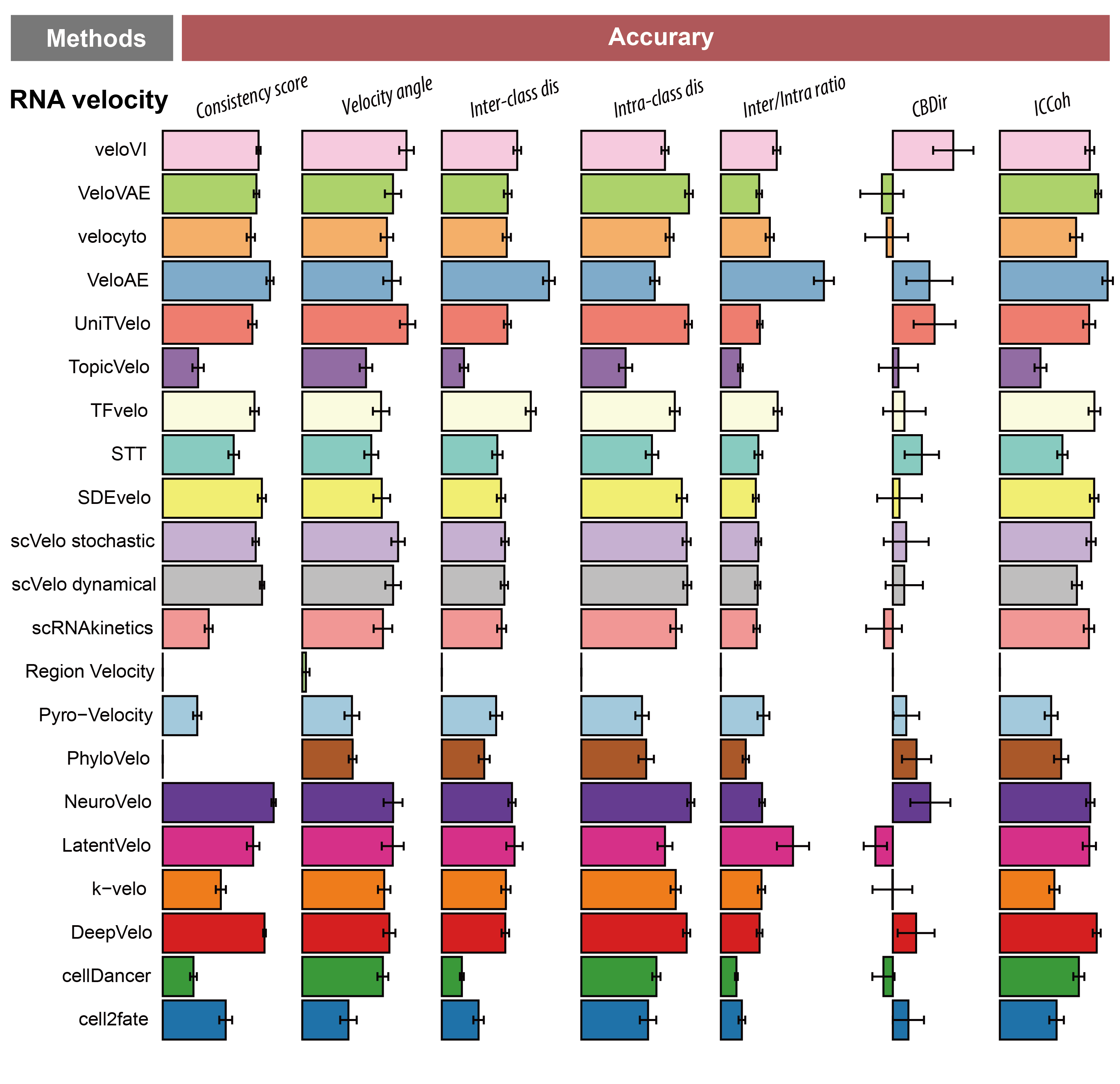

### Supplementary Figure 2.png

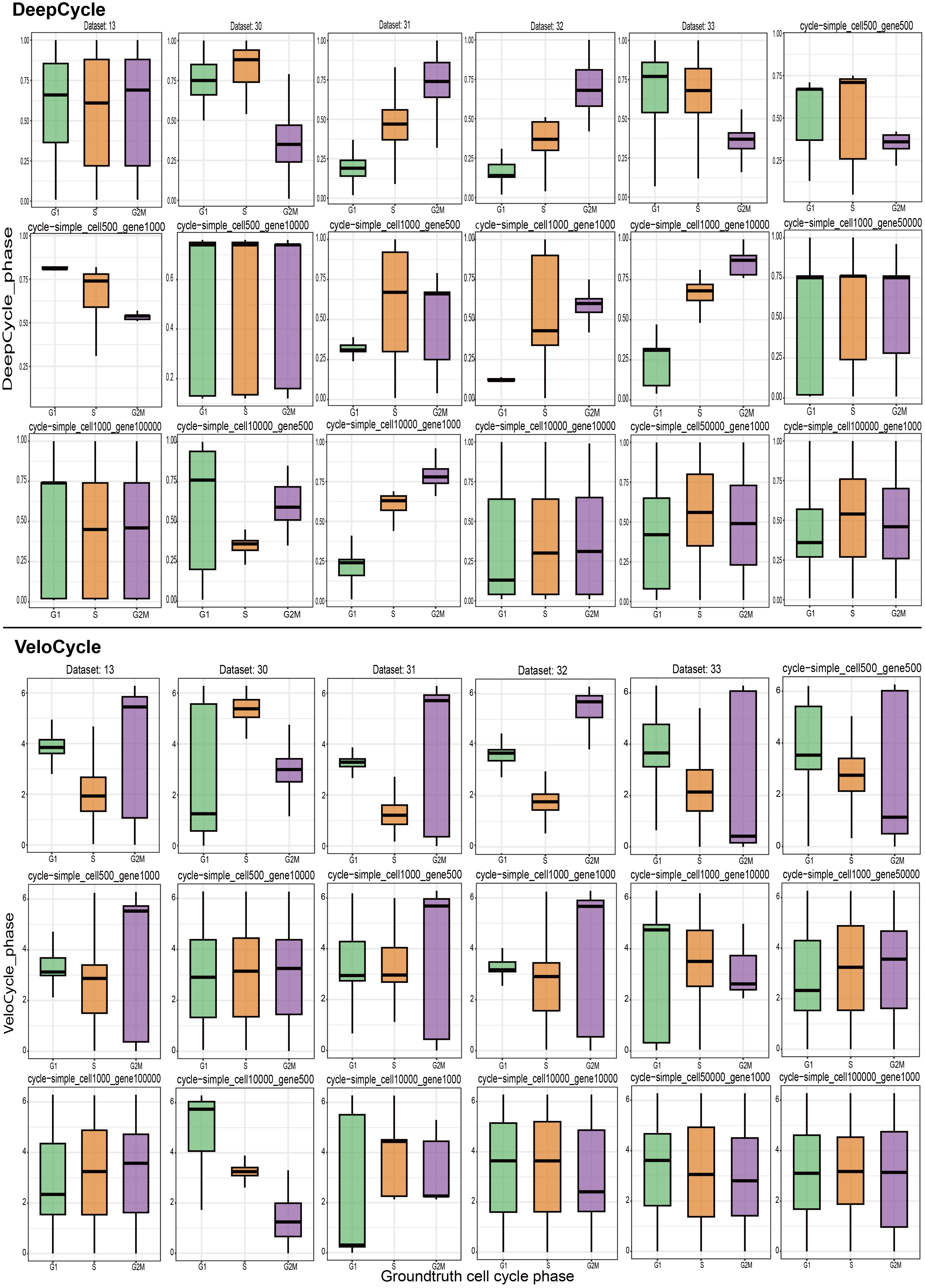

### Supplementary Figure 3.png

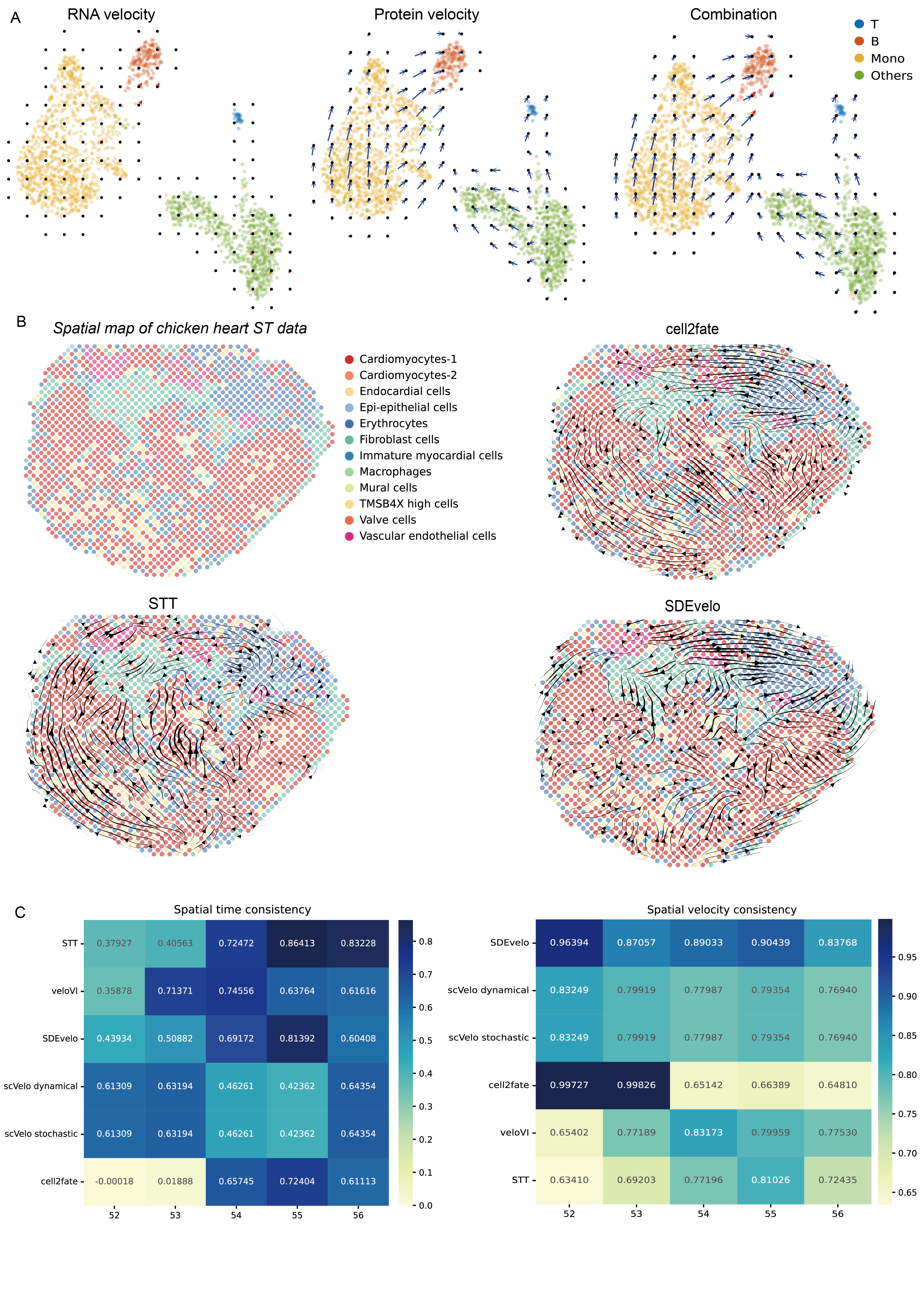
